# Considerations for Deconvolution: A Case Study with GTEx Coronary Artery Tissues

**DOI:** 10.1101/2022.05.17.492324

**Authors:** Zachary P. Brehm, Valeriia Sherina, Avi Z. Rosenberg, Marc K. Halushka, Matthew N. McCall

## Abstract

Differential expression analyses are ubiquitous in the realm of statistical genomics, used to estimate functional differences between genomes of groups of subjects. However, differences in tissue composition between groups may contribute to changes in gene expression, potentially obscuring the detection of functionally significant genes of interest. Deconvolution techniques allow researchers to estimate the abundance of each cell type assumed to be in a tissue. While deconvolution is a useful tool to estimate composition, several crucial considerations must be made when setting up and employing such a workflow in an analysis. We perform a deconvolution on GTEx coronary artery data using CIBERSORT and discuss the challenges and limitations in order to highlight future areas of improvement in the deconvolution framework.

## Introduction

Whole tissue sequencing measures the gene expression of a diverse array of cell types which may exist in a variety of states. Tissue composition is an influential factor in the result of bulk RNA-seq experiments on whole tissues. Often in such studies we are interested in studying functional differences between experimental groups. However, if we do not account for tissue composition, we may confound these effects due to differences in form (1–8).

Deconvolution methods have been developed to identify the compositional abundance of cell types present in complex tissues. The most common methods either rely on cell reference profiles or cell type specific markers for component cell types. CIBERSORT (9) is a popular method, which uses cell reference profiles to estimate cell type proportions in whole blood expression data. Subsequently, this method has been extended to tissue samples; however, reference profiles are not precomputed for tissue data and must be estimated using user-supplied cell type data. These reference profiles are used to identify genes that distinguish between cell types and are used to perform a deconvolution of tissue expression data to identify the relative abundance of each cell type in the tissue. In contrast, reference-free methods, such as TOAST (10, 11), rely on sets of marker genes for each cell type to identify the composition of complex tissues.

Single cell RNA-seq technology provides an alternative method to identify functional differences between specific cell types. Methods based on single cell RNA-seq, such as MuSiC (12), seek to identify the cell type specific expression profile of cell types in this way. While these methods are effective, single cell data does not necessarily describe the gene expression of cells as they exist in a human subject. The method of isolating single cells can potentially influence gene expression. For example, culture effects are a potential hazard of growing cells in vitro, while harvesting cells from tissue may damage them, altering expression.

The Genotype Tissue Expression (GTEx) study was designed to identify expression quantitative trait loci (eQTLs) in 54 normal human tissues using paired DNA and RNA-sequencing data from rapid autopsies of >900 individuals (13). Additional information collected as part of the GTEx study includes digitized images of adjacent tissue, along with rich phenotypic and technical variables. GTEx has been enormously successful in meeting its objectives while providing a rich resource for ancillary studies (14–21).

Among the tissues collected by GTEx, coronary artery tissue is an obvious target for the application of deconvolution methods. In healthy subjects, these tissues consist of a variety of cell types that vary in their abundance between tissue samples. Additionally, cardiovascular disease often manifests in the coronary artery in the form of an atherosclerotic plaque. Atherosclerosis disease progression is characterized by the formation and potential growth of an eccentric plaque composed of variable amounts of inflammatory cells, lipids, cholesterol crystals, calcification, extracellular matrix material, and mesenchymal cells (22). These compositional changes may obscure functional changes within component cell types that are also associated with disease progression.

While deconvolution methods are useful to gain insight into the composition of whole tissue sequencing data, there are several crucial considerations that must be made during their use and further application in downstream analyses. In this paper we highlight some of the challenges that investigators should consider when applying deconvolution methods to whole tissue sequencing data and using the composition estimates in downstream analyses. We use the analysis of coronary artery RNA-seq data to demonstrate these challenges and suggest potential paths forward.

## Methods

### Obtaining GTEx data

The Genotype Tissue Expression (GTEx) study provides histological images and RNA-seq data for hundreds of coronary artery tissue samples. These samples include a diverse population of atherosclerotic tissues. To access the histological images for these samples, we used the histology viewer webtool on the GTEx Portal. We then obtained the corresponding RNA sequencing data using the recount3 Bioconductor package (23, 24). Coronary artery samples were obtained by first downloading the “BLOOD_VESSEL” labeled data as a SummarizedExperiment object and then selecting samples labeled as “Artery – Coronary” in the gtex.smtsd field of the column data.

### Obtaining SRA data

Deconvolution with CIBERSORT requires a set of reference expression profiles for each of the cell types assumed to be in the tissue. We also use the recount3 package to access reference samples from the Sequence Read Archive (SRA) for relevant cell types that were initially identified using the MetaSRA (25) tool by searching for cell types found in coronary arteries: smooth muscle, endothelial, macrophage, lymphocyte, erythrocyte, plasma, serum, foam cells, adipocytes and cardiac muscle cells. The SRP numbers associated with the search results from MetaSRA for each cell type were used to access these samples through recount3.

### Processing GTEx data

The GTEx coronary artery data provided by recount3 contains 253 RNA sequencing samples of 63,856 transcripts. These data were filtered to ensure that all samples were obtained from unique subjects. Five of these samples were from duplicated subjects and were removed. We filtered the number of features by removing all transcripts from which we observed counts less than 1,000 in all samples. After this filtering step, 12,642 transcripts remained. The counts were transformed according to the scaling procedure employed by the transform_counts function of the recount3 package.

### Processing SRA data

The initial set of cell type reference data was filtered to remove samples with <1 million total reads and those mischaracterized by cell type. An exploratory analysis suggested that many samples did not match the cell type described by the search results. After consulting the metadata on SRA itself, we corrected the cell types when applicable or eliminated samples that did not match any of the desired cell types. We then batch corrected using the RUVr function from the RUVSeq package (26) with *k* = 68.

### Quantification of atherosclerosis in GTEx coronary artery histopathology

Each GTEx coronary artery sequencing dataset is paired with a histology image depicting generally two segments of tissue. The digitized images were scored for atherosclerotic plaque type using the scheme of Virmani et al. (22) by a cardiovascular pathologist. Fig. 1 provides examples of the many of the different plaque scores across the tissues. In addition to the plaque score, each sample was scored for the degree of inflammatory cells: 0 No inflammation; 1-Any inflammatory cells; 2 -Heavy Inflammation, adipose tissue: 0 - No Fat (*<* 10% of the wall); 1 - Slight Fat (10 − 25% of the vessel wall); 2 - Moderate Fat (25 − 100% of the vessel wall); 3 - Extensive Fat (*>* 100% of the vessel size), and myocardium tissue: present or absent in the histology. If the plaque scores of the different pieces of tissue in an image disagreed, the maximum of the two scores was assigned to the sample.

**Fig. 1.**
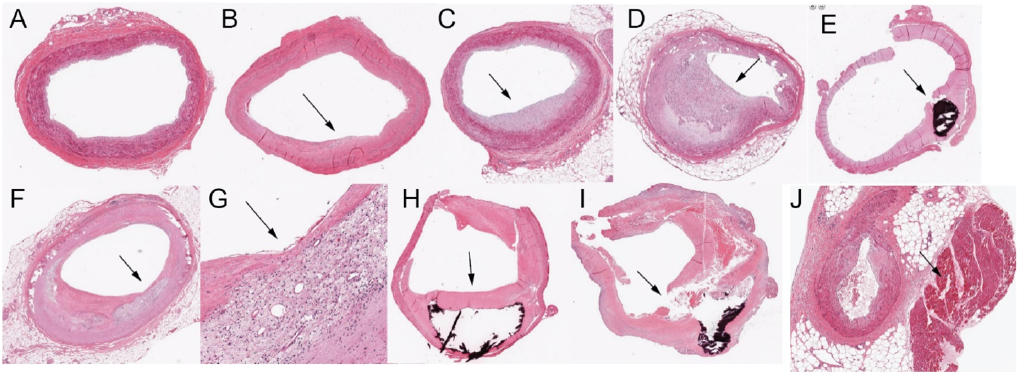
Atherosclerotic plaque types among the GTEx coronary artery samples. Arrows point to the histopathology or unexpected finding. A) normal coronary/intimal thickening; B) intimal xanthoma C) pathological intimal thickening D) fibrous cap atheroma E) calcified nodule F) thin fibrous cap atheroma G) high power view of the thin fibrous cap H) fibrocalcific plaque I) plaque rupture (with hemorrhage into atheroma) J) adjacent myocardium. All images from the GTEx tissue collection.

### Identifying high confidence samples with both histopathology and RNA-seq

The presence of two pieces of tissue per histology image, often with different plaque patterns, precluded certainty as to which piece best represented the gene expression data. To resolve this, we selected samples according to a combination of plaque score and a constructed macrophage score. Macrophage expression levels were determined by summing the expression counts in each sample across the macrophage marker genes listed in Table 1. These sums were then centered by their mean and scaled by their standard deviation to form a pseudo z-score.

**Table 1.** Marker genes used to define smooth muscle cells (SMC), endothelial cells (END), erythrocytes (RBC), adipocytes (ADI), cardiomyocytes (CMC), lymphocytes (LYM), and macrophage (MAC) cell types. These genes were obtained through a combination of the literature on single-cell studies of atherosclerosis (32) and confirmed for specificity using the Tabula Sapiens reference in the cellxgene webtool (33).

| CELL TYPE | GENE |
| --- | --- |
| SMC | ACTA2, TAGLN, TPM1, MYH11, NOTCH3, MYL9, MYLK, TPM2 |
| END | PECAM1, VWF, CD34, TIE1 |
| RBC | HBA1, HBA2, HBB |
| ADI | ADIPOQ, ADIPOQ-AS1, LEP |
| CMC | MYH6, MYH7, MYL3, MYL4, MYL7, MYLK3, MYBPC3, ACTN2, TTN, TNNT2, XIRP1 |
| LYM | CD2, CD3D, CD4, CD8A, CD19, CD22, CD69, CD79A, CD79B, BANK1, BCL11A, JCHAIN, CCL5, GZMK, FGFBP2, PRF1, ADGRG1, SPOCK2, IL32, IL7R, TRBC1, TRBC2, TRAC |
| MAC | C1QA, C1QB, C1QC, MSR1, APOC1, APOE |

Let *y*_*ij*_ denote the count of marker gene *i* from subject *j*. For each cell type *k ∈ {* 1, 2, …, *K}*, let 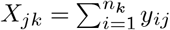 where *n*_*k*_ denotes the number of marker genes for cell type *k*. Let 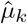 and 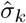 denote the mean and standard deviation of *X*_*jk*_ for each *k*, respectively. We then define the cell type score of cell type *k* by

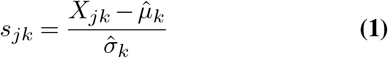

The control group is defined as the samples with a plaque score of zero and a macrophage score below the median value, while the atherosclerosis group is defined as the samples with a non-zero plaque score and a macrophage score above the median value.

### Estimating composition of GTEx coronary artery

Composition estimates for each of the cell types assumed to be in the coronary artery samples were estimated using CIBERSORT. GTEx coronary artery samples and our curated reference cell type data were uploaded to the CIBERSORT web tool. CIBERSORT was run with a kappa of 999, q value of 0.3, a minimum of 50 and maximum of 150 barcode genes per cell type, and without the quantile normalization step.

### Power analysis for RNA-seq data

Power analyses were performed with the PROPER method (27). This method assesses the marginal power of a study to detect differentially expressed genes according to several prespecified datasets with a diverse range of inherent biological variability. Of these prespecified datasets, we used the one with the greatest amount of biological variability between samples. These analyses were performed with 20 simulations for each set of conditions, where the data is simulated to have 1, 5, or 10% of genes be differentially expressed.

### Composition adjusted differential expression analysis

To perform a composition-corrected differential expression analysis on the GTEx coronary artery data, we utilized the DESeq2 package (28). To account for the composition of the tissues, we considered two models: (1) a model that used CIBERSORT estimates of composition or (2) a model that used pseudo z-scores defined by Eq. S.

To place the CIBERSORT estimates on a similar scale to those obtained through the gene expression itself, we centered the estimates by their mean and scaled them according to their standard deviation as we did with the expression-based estimates. These estimates were then included as covariates in the DESeq2 model, except for the macrophage and lymphocyte estimates due to their strong correlation with the plaque scores used to define the groups of interest. Additionally, CIBERSORT estimates for erythrocyte composition of the artery samples were not useable in this framework as they were essentially all estimated to be zero. Thus, we included the expression-based estimates for erythrocyte composition in both DESeq2 models. Additional covariates in the models were the subjects’ sex, age, and the year during which a sample was sequenced. Age is an important factor in these samples as older subjects have had more time for atherosclerotic plaques to develop in the artery, while sequencing year was included to account for technical variation introduced by sequencing these samples from 2012 to 2017.

## Results

### Processing GTEx data

In our initial exploratory analyses, we sought to identify any subpopulations of interest in the GTEx data. We performed a principal components analysis to first check for any dominating factors that would influence the downstream analyses. Supplementary Fig. 1 shows the first two components of this PCA after selecting the top 100 most variable genes in terms of absolute fold change between groups of high and low plaque samples in the GTEx data. The second component separates the sex of the subjects in the GTEx data. The primary genes associated with this split were Y chromosome genes and XIST. This motivated the need to include the subjects’ sex as a covariate in the downstream differential expression models. After sex correction, we found a small group of samples that clustered together in a t-distributed Stochastic Neighbor Embedding (t-SNE) (29– 31) dimension reduction plot shown in Supplementary Fig. 2. Histologic review of these samples indicated these were cardiac vein samples based on the presence of at least one of the two tissues being venous. As we were only interested in coronary artery tissue, these 12 samples were removed, leaving 235 for downstream analyses.

### Processing SRA reference data

The MetaSRA search identified 5,188 samples. An exploratory analysis of clustering in a multidimensional scale reduction plot showed that many samples did not cluster with their designated cell types. A manual check of the metadata for each sample found a substantial number of discrepancies between the cell types reported by MetaSRA and the cell types recorded in SRA. Hand curation validated with batch correction using RUVSeq resulted in a final reference set of 790 samples (Fig. 2). These samples represent 7 main cell types: smooth muscle cells (SMC), endothelial cells (END), erythrocytes (RBC), lymphocytes (LYM), macrophages (MAC), adipocytes (ADI), and cardiomyocytes (CMC). These data were used as reference data to deconvolute the coronary artery samples using CIBERSORT. We similarly filtered the transcripts found in the reference data by removing all transcripts with counts less than 1,000 in all samples. The filtered reference data contains 19,329 transcripts that passed this condition.

**Fig. 2.**
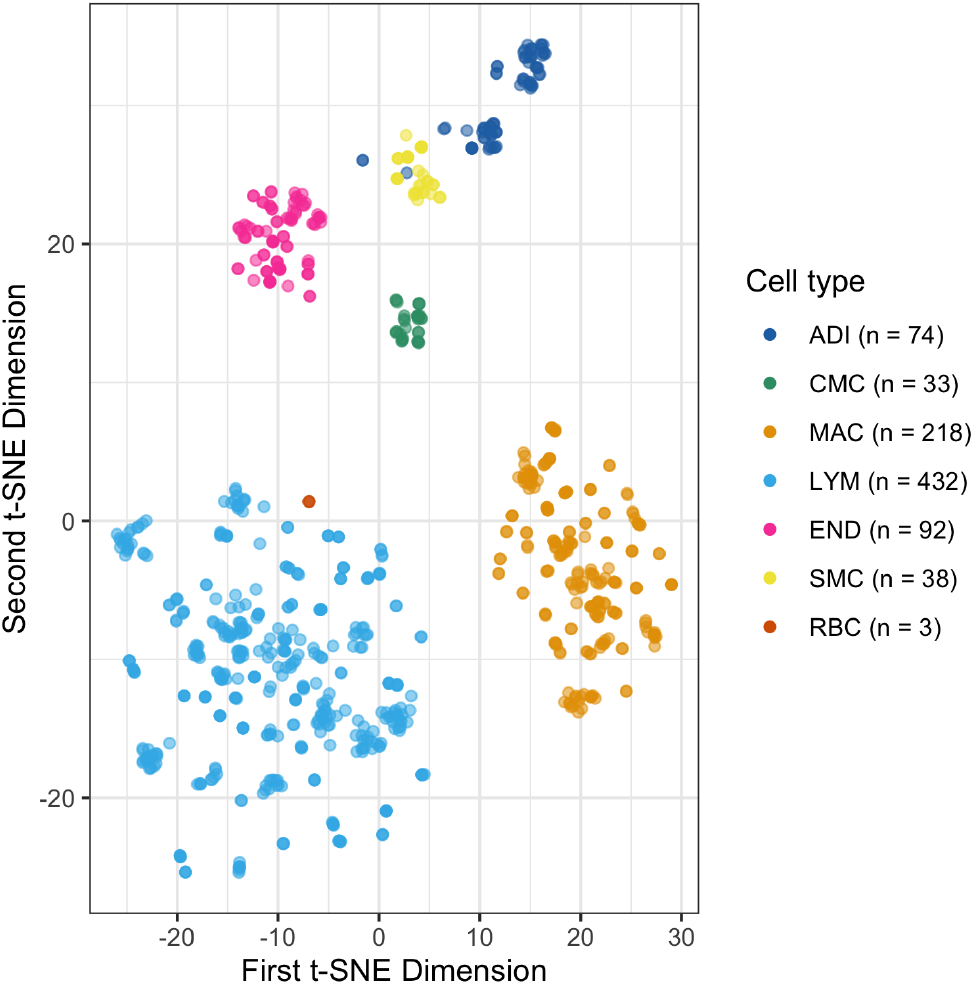
Finalized reference data set clusters by cell type following t-SNE. The cell types shown are smooth muscle cells (SMC), endothelial cells (END), erythrocytes (RBC), lymphocytes (LYM), macrophages (MAC), adipocytes (ADI), and cardiomyocytes (CMC). Dimension reduction was performed using the Rtsne package with perplexity of 40.

### Identifying high confidence samples with both histopathology and RNA-seq

For each GTEx coronary artery sample that underwent RNA sequencing, there are often histological images of two segments of coronary artery. The plaque scores for these two segments often differed, complicating their use in a differential expression analysis to detect gene expression changes associated with plaque score. The presence of macrophages is highly correlated with atherosclerotic plaques, thus samples with advanced atherosclerosis should have elevated expression of macrophage genes. Therefore, we selected the 138 samples with agreement between atherosclerosis score and expression of macrophage gene markers. Of these 138 high confidence samples, 53 were deemed free of atherosclerosis (0 plaque score and below median macrophage expression), and 85 were identified as atherosclerotic (non-zero plaque score and above median macrophage expression).

We then explored cell composition expression patterns across the GTEx samples (Fig. 3) using known cell type specific genes (Table 1). Cardiac muscle cells and red blood cells identify individual samples noted by generally consistent expression across samples with few outliers expressing abnormally high cardiac muscle genes. Likewise, there are numerous samples with elevated expression of adipose tissue. Note that the distribution of cardiac muscle cells, red blood cells, and adipocytes are unrelated to plaque or macrophage score. Conversely, genes from lymphocytes and macrophages have little relative expression in plaque type 0, while the samples with the highest expression are typically found in more severe plaque types. For lymphocytes, marker genes of T-cells are relatively higher than those of B-cells, which have more of a binary off/on pattern like cardiac muscle and erythrocytes. There appears to be less similarity between the samples partitioned according to histologic plaque type than expected, consistent with the observed discrepancy between gene expression data and adjacent histology.

**Fig. 3.**
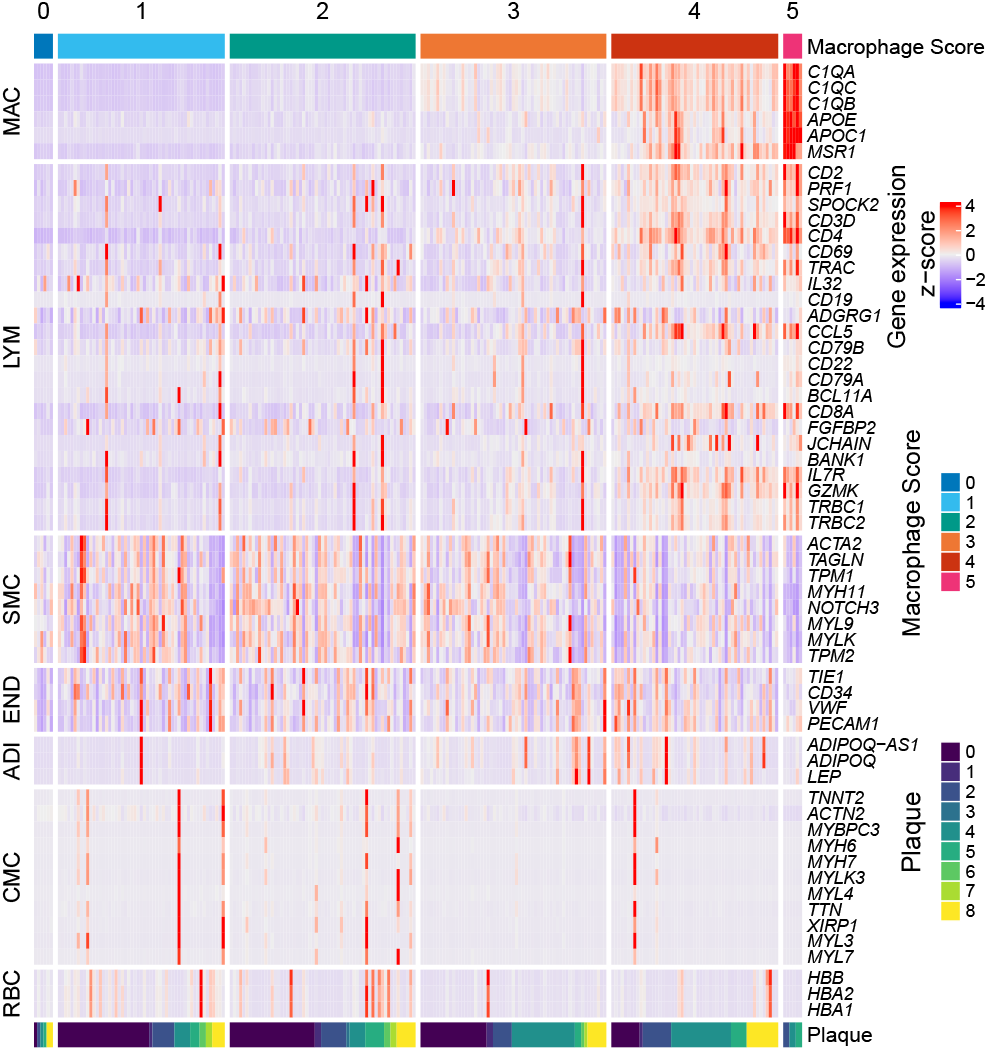
Difference in composition between tissues drives variance in expression. Expression of marker genes in GTEx coronary artery for macrophage (MAC), lymphocyte (LYM), smooth muscle cell (SMC), endothelial cell (END), adipocyte (ADI), cardiomyocyte (CMC), and red blood cells (RBC). Macrophage scores correspond to the quantiles of the macrophage expression score described by equation 1. (0 – [0, 2.5]; 1 – (2.5, 25]; 2 – (25, 50]; 3 – (50, 75]; 4 – (75, 97.5]; 5 – (97.5, 100])

### Estimating composition of GTEx coronary artery

We used CIBERSORT to estimate the composition of each coronary artery tissue (Fig. 4). As expected, smooth muscle cells were the predominant cell type in most samples, with endothelial cells as second most abundant cell type. Macrophage estimates were greater than lymphocyte estimates, consistent with their known higher abundance in atherosclerotic plaques. Adipocyte cell signals were widely variable across samples, indicating variation in the amount of adventitial tissue. Notably, this signal did not correspond well to histology-based estimates of the proportion of adipose Supplementary (Fig. 3). Despite general consistencies with expected results, we highlight some concerns with these estimates.

**Fig. 4.**
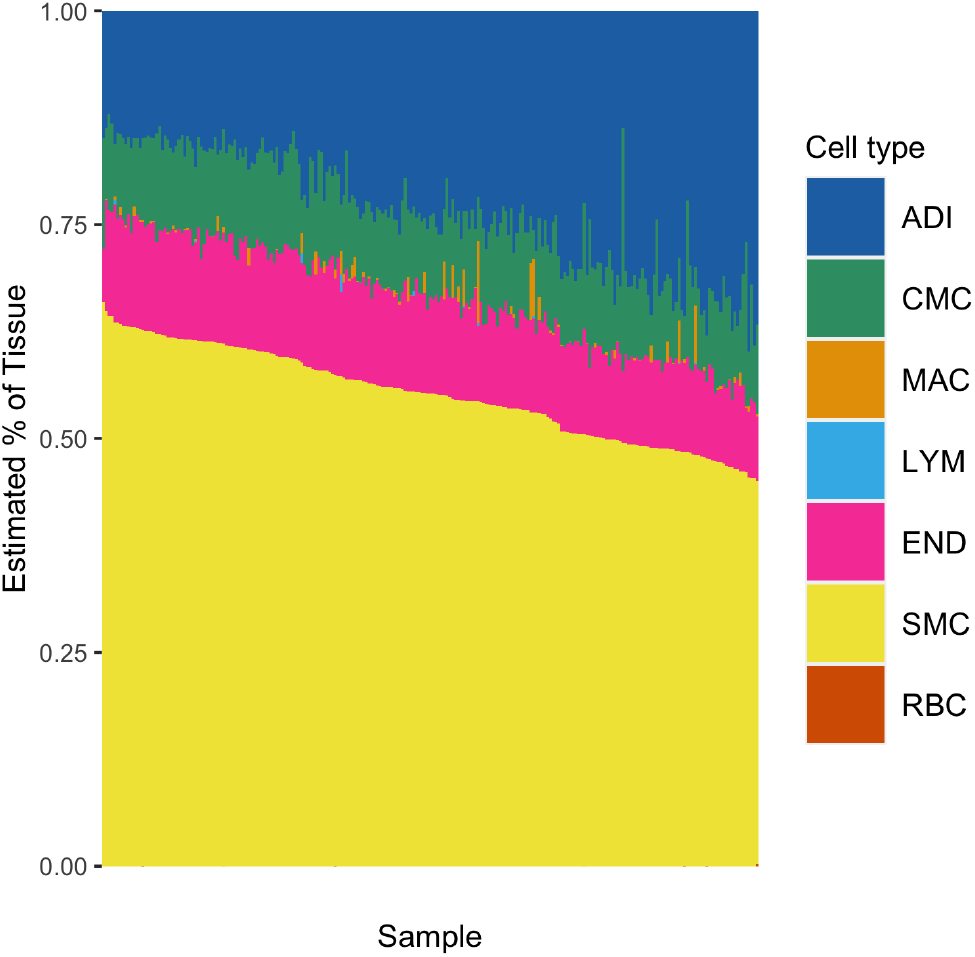
Deconvolution of GTEx data with CIBERSORT. Composition estimates of macrophage (MAC), lymphocyte (LYM), smooth muscle cell (SMC), endothelial cell (END), adipocyte (ADI), cardiomyocyte (CMC), and red blood cells (RBC) in GTEx coronary artery. Each strip on the x-axis is one sample from GTEx, and each strip describes the percentages of each cell type in the sample.

First, as noted in Fig. 3, cardiac muscle is rarely seen, yet from CIBERSORT, muscle cells appeared as generally expressed in all samples. The histological assessment does not support the presence of cardiac muscle tissue with such high frequency. We believe this occurs because of similarities that smooth muscle and cardiac muscle cells have as muscle tissue, leading to some correlation of genes for these cell types in the CIBERSORT signature matrix. The range of these estimates seems appropriate; however, more mass in their distribution should be shifted towards 0. Adipose tissue estimates appear robust at the lower boundary of estimates but appear overexpressed at the upper boundary region. While adipose tissue can occupy a large amount of area in these samples, adipocytes have a relatively small amount of RNA per unit area due to their structure. Erythrocytes are almost always given a proportion estimate of 0. However, Fig. 3 clearly demonstrates numerous samples with substantial expression of RBC-specific hemoglobin genes: HBA1, HBA2 and HBB. Finally, lymphocytes are underrepresented in these estimates. From Fig. 3, it is clear this cell type is expressed by many samples and generally in strong correlation with macrophage expression.

Fig. 5 demonstrates the relationship between the composition estimates obtained from CIBERSORT and the composition estimates derived from the expression of marker genes for the cell types found in the coronary artery tissue. Due to a large proportion of zeros, the RBC estimates were omitted from this Figure and instead are shown in Supplementary Fig. 5. The CIBERSORT estimates of lymphocyte composition fail to capture the variation described by the expression scores in the sample. In contrast, cardiac muscle estimates exhibit too much variance that is not present in the expression data. Smooth muscle cell estimates with largely average expression have a considerable amount of variability within the CIBERSORT estimates. Finally, there is a negative correlation between the endothelial cell estimates from CIBERSORT and the endothelial cell expression scores.

**Fig. 5.**
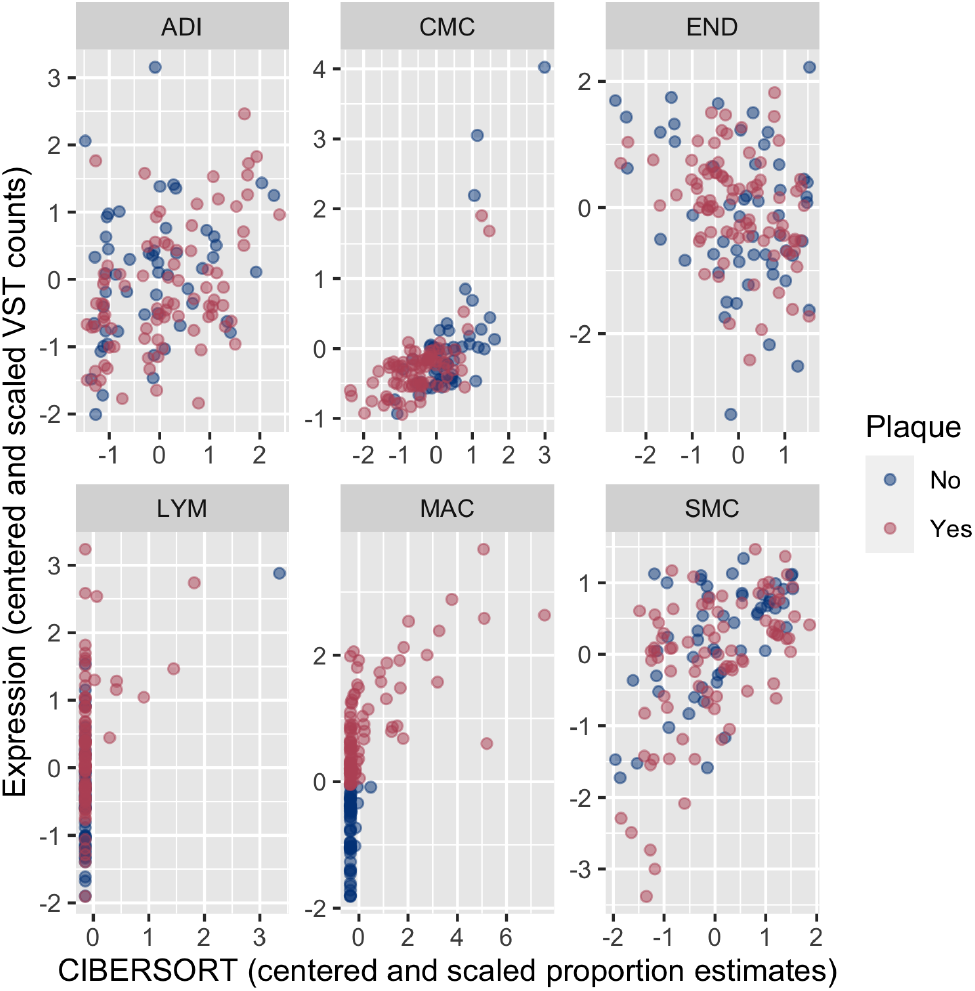
Comparison of CIBERSORT composition estimates with gene expression derived composition estimates.

**Fig. 6.**
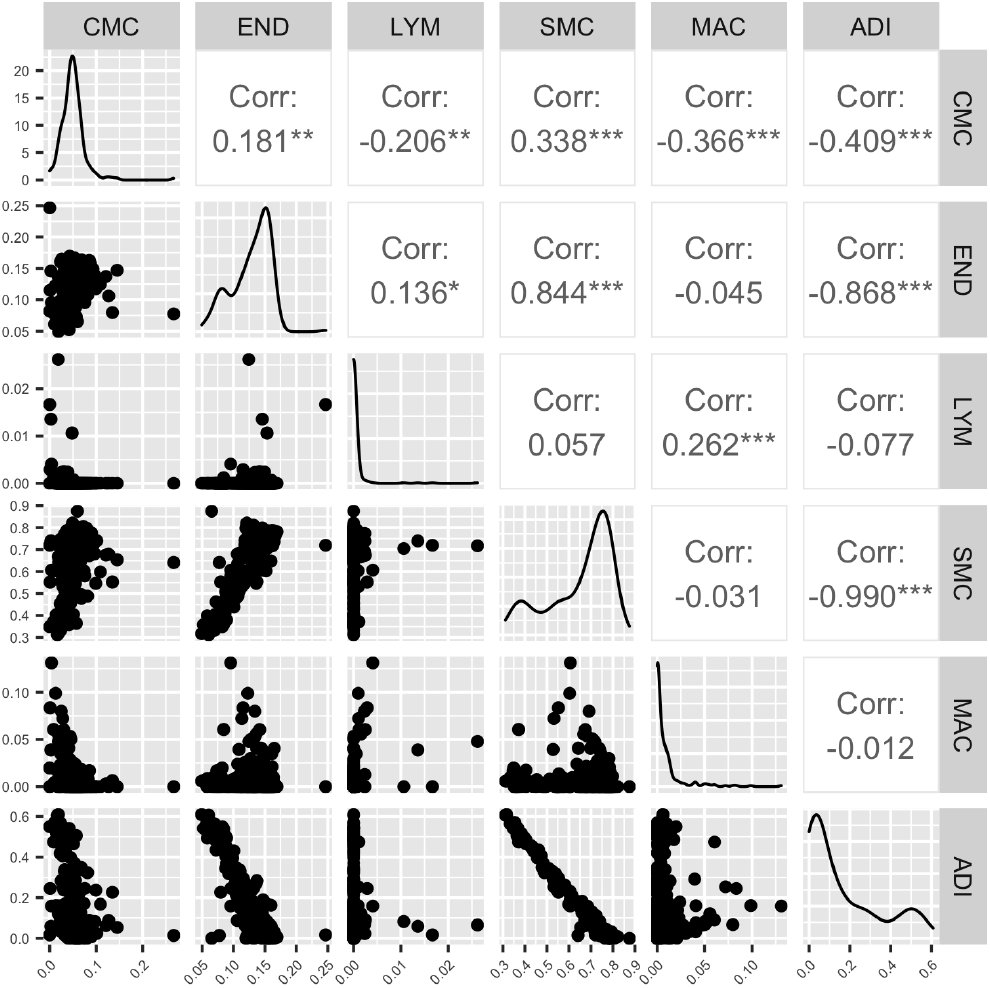
Correlation between cell type composition estimates. Correlation of composition estimates from CIBERSORT between macrophage (MAC), lymphocyte (LYM), smooth muscle cell (SMC), endothelial cell (END), adipocyte (ADI), cardiomyocyte (CMC), and red blood cells (RBC) in GTEx coronary artery.

A deconvolution of tissue gene expression via CIBERSORT requires an initial list of cell types assumed to be in the tissue; however, there are no well accepted methods for determining the list of cell types needed to use for deconvolution. Adjacent histological images can be used to identify cell types present in tissues; however, this is resource intensive and subject to user bias. To demonstrate the effect of component cell type selection, we compared CIBERSORT estimates including or excluding the red blood cell reference samples. CIBERSORT estimated that all GTEx samples lacked erythrocyte composition, suggesting that RBCs were an extraneous cell type that did not need to be accounted for in the mixture. Therefore, the results should largely be the same whether RBCs were included in the deconvolution or not. However, the CIBERSORT composition estimates were dramatically different depending on whether RBCs were included in the model (Fig. 7). Most importantly, there was a substantial reduction in the estimated contribution of smooth muscle cells to the tissue when red blood cells were not included in the deconvolution, with the median estimate falling from 70.7% to 27.8%. The smooth muscle cell signature is generally the largest of all the cell types in the initial results, but it becomes largely outweighed by the endothelial cell estimates which see an increase in median estimated composition from 13.5% to 52.9%. Previously, and like the erythrocytes in question, lymphocytes were largely estimated to have no presence in the tissue composition, with the largest estimated percentage to be 2.6%. In the updated estimates, the median contribution is still 0%, however the maximum has jumped from 2.6% to 62.8%. Cardiac muscle cell composition was similarly affected, seeing a relatively stable median composition estimate of 4.8% holding throughout, but a maximum estimate jumping substantially from 26.7% to 86.2%. These differences highlight how significantly the composition estimates from CIBERSORT can change and the need for a rigorous assessment of the cell types assumed to be present in tissues that we wish to deconvolute.

**Fig. 7.**
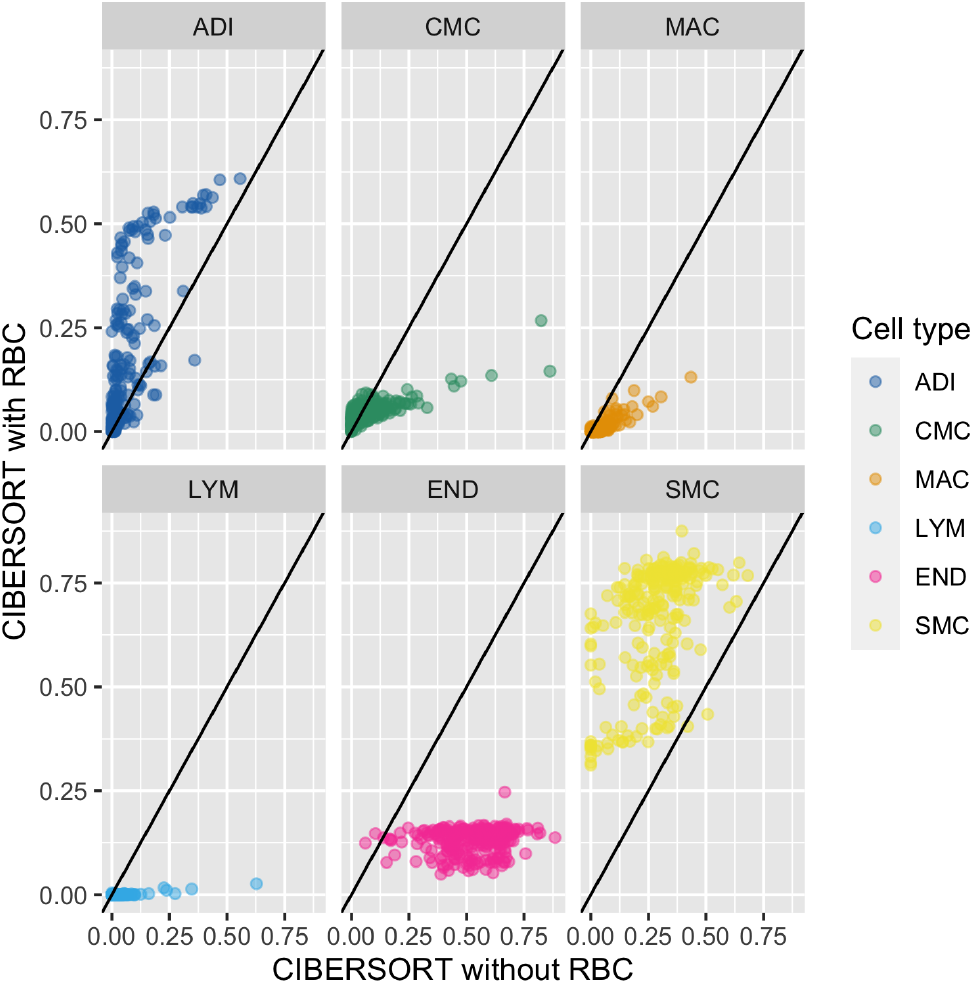
Shift in CIBERSORT composition estimates when erythrocyte cells are not included in the deconvolution.

### Power analysis for RNA-seq data

Power analyses were completed using the PROPER package to simulate experiments under several conditions. We consider cases when 1, 5, and 10% of 12,642 transcripts are differentially expressed with a log2 fold change (l2fc) 0.585 or 1. Each condition considers two groups of subjects with 53 and 85 samples in each group, as in our GTEx data, and a nominal false discovery rate (FDR) of 0.1. Of the 6 different combinations of parameters, 5 of the simulation results met the desired power of 0.8, while the one failing to meet this condition had a marginal power of 0.779. The full results are shown in table 2.

**Table 2.**
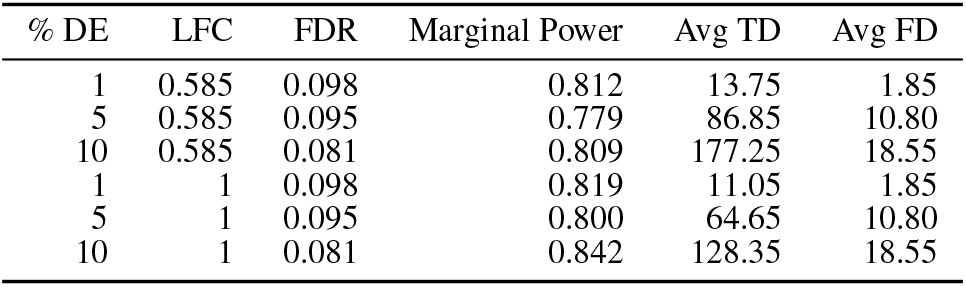
Power analysis results from PROPER.

### Composition adjusted differential expression analysis

We performed a DESeq2 analysis with two separate sets of composition estimates: (1) those obtained from CIBERSORT and (2) those obtained by generating pseudo z-scores directly from the expression data. We additionally performed a DESeq2 analysis with no composition correction to account for a composition agnostic approach. After removing any transcripts with a base mean below 500, we filtered the results tables to include transcripts with an adjusted p-value below 0.1. The CIBERSORT adjusted model was most conservative, finding 2,180 significant transcripts, while the expression adjusted model found 3,575. The model which did not adjust for composition found 4,528.

Supplementary Fig. 6 displays the distribution of log2 fold change estimates from the DESeq2 results in each model. Of the transcripts mentioned above, we selected the top 400 in absolute fold change that were significant in each model and then took a selection of 100 transcripts from the cumulative list. The general effect of including composition estimates within the DESeq2 model results in a shrinkage towards zero on the fold changes observed in the results.

We must note that each of the methods used to estimate composition are dependent on the GTEx data we are analyzing with DESeq2. This dependence between the composition covariates and data inflates the false discovery rate, making any inference on these results problematic.

## Discussion

In this paper we have performed a complete deconvolution workflow with CIBERSORT on the GTEx coronary artery data. We have found that while deconvolution methods are useful tools for estimating the composition of tissues from sequencing data, there are numerous considerations to be made when undertaking such an analysis that have a profound effect on the composition estimates and gene expression differences. In particular, we highlight the difficulties encountered while working with public data to prepare for, undertake, and assess a deconvolution workflow using CIBERSORT and DESeq2.

### Preparing the data

We have worked with two sets of data in this paper, the SRA reference set of cell type specific expression counts and the GTEx coronary artery set of whole tissue expression counts. Both sets of data featured specific challenges.

When acquiring public data, there are many repositories and processing techniques that may be employed to process samples. Metadata collection and reporting is rarely aligned between different experiments, and identifying samples of interest can be particularly challenging when reviewing these data to ensure accuracy and completeness of sample information. Additionally, batch correction methods have long been used to account for technical artifacts between experiments; however, when analyzing publicly available cell type data, often all the data from a given experiment comes from the same cell type, potentially confounding batch and cell types. Adjacent histology for sequencing data can be an effective tool for assessing the potential composition of tissue samples, however these images are not without limitations. As we have seen with the GTEx data, if images contain multiple examples of tissue available for sequencing, it can be difficult to determine which piece is closest to the sequenced sample. In our case particularly, the degree of atherosclerosis in the sequencing data may be difficult to determine if the plaque types between two samples are substantially different. Other histological factors may aide in determining which piece is the likely candidate.

Further, by histology, some samples labeled as coronary arteries were in fact veins. These samples also showed gene expression patterns that were distinct from the true artery samples (Supplementary Fig. 2), indicating another cause for concern when using GTEx data (17).

### Performing a deconvolution with CIBERSORT

Perhaps the most critical step of the deconvolution process is determining which cell types should be assumed present in the tissues. As we have shown in this work, small and seemingly inconsequential changes to the set of possible cell types can result in substantial shifts in the distribution of estimates that we observe, as with the red blood cell example. How distinct do the different cell types of consideration need to be? Here we use fairly broad classifications of cell types, grouping all lymphocytes together. In this case, such cell types are considered different enough from others like smooth muscle cells that this should be sufficient. However, if there is interest in differences between immune cell composition or activity, one may wish to further separate lymphocytes into B cells and T cells or even finer subsets. Additionally, similarities between cardiomyocyte and smooth muscle cell gene expression appears to have made it difficult for CIBERSORT to assign a zero value to the estimates for cardiomyocytes, as suggested by histology and marker gene expression Fig. 3.

### Employing composition in analyses

With composition estimates, we must consider that there are different effects of correlation to account for. We can broadly group the causes of these correlations into those of biological origin and technical origin. Correlations that are biological in nature result from cases like similar biological function, e.g. immune cells, muscle cells, etc. A technical source of correlation is introduced by the sum to one constraint of the composition estimates. With atherosclerosis in particular, macrophages are so strongly correlated with the condition of interest and lymphocytes are so strongly correlated with macrophages that the inclusion of either of these two cell types in the DESeq2 analysis is suspect.

Finally, inferential procedures that utilize cell type composition estimates as we have obtained them pose a challenge. The estimates that we have obtained are based on the GTEx data itself, and as such they are effectively present on either side of the regression equation that differential expression methods are based on, increasing the type I error rate (34).

### Conclusion

We have described several challenges in any deconvolution-based analysis of differential expression through a case-study of atherosclerotic vessels among GTEx data. Not all cell specific expression datasets are properly annotated and the use of these samples requires additional validation. Histological images in GTEx or other tissue collections do not faithfully indicate the same composition as the material collected for RNA expression studies. CIBER-SORT is particularly sensitive to the cell types included in the matrix, where even having a cell type (RBC) that showed no expression profoundly affected composition estimates. In conclusion, we expect this study will raise awareness of some specific considerations when performing tissue deconvolution analyses.

## ACKNOWLEDGEMENTS

National Institutes of Health (R01HL137811 and R01GM130564)

## Supplementary Information

### Supplementary Note 1: Subject sex driver of expression diversity in GTEx data

After selecting for the top 100 genes in absolute log fold change, we found that the sex of the subjects in the GTEx coronary artery data was a primary driver of separation along the second principal component of the data.

**Supplementary Figure 1.**
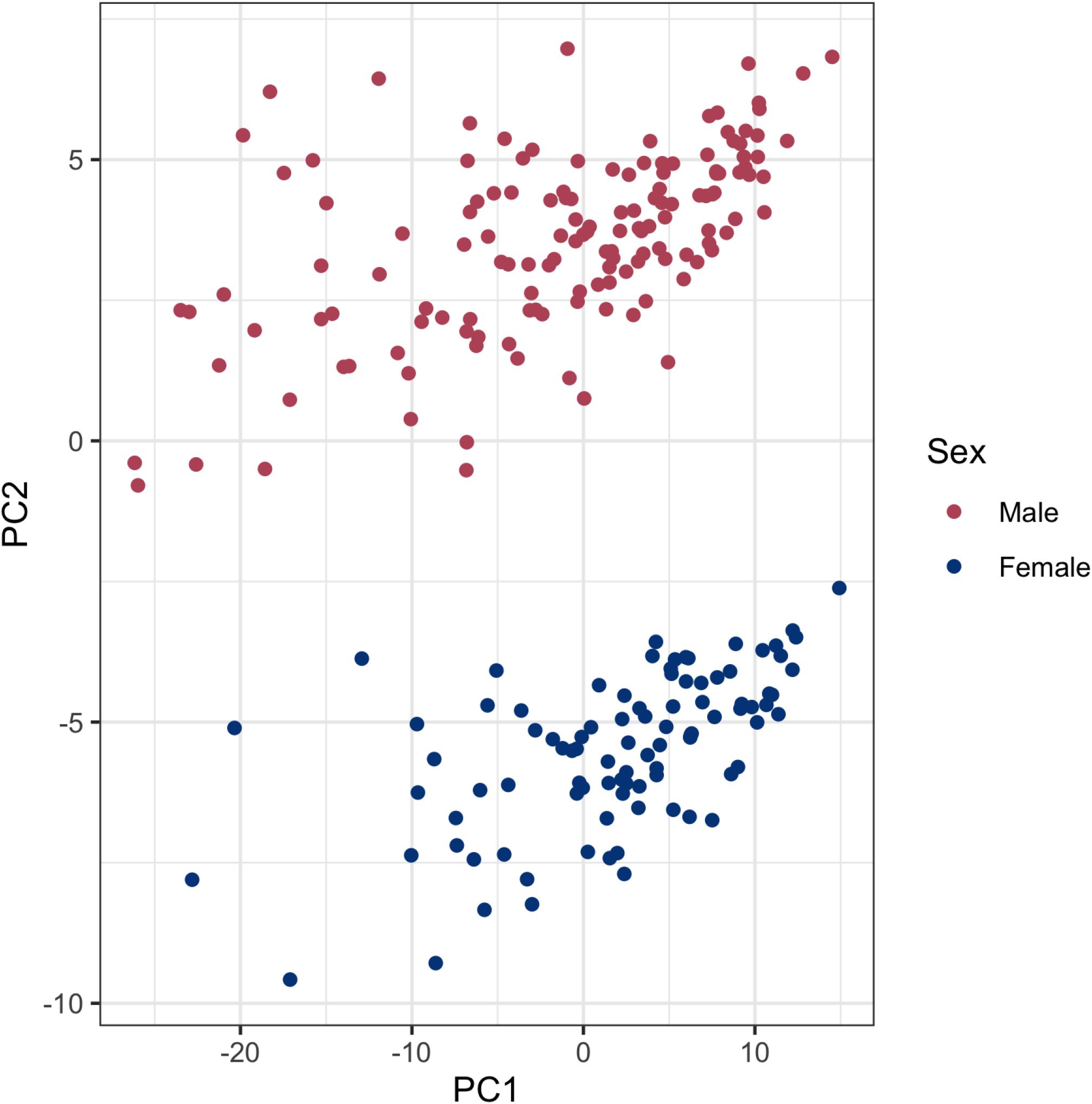
Initial PCA attributes largest differences to subjects’ sex.

### Supplementary Note 2: Vein tissue among coronary artery samples

During exploratory analysis we discovered an outlier cluster of GTEx samples which are likely vein tissue. From a prior review of the histology, we knew that several pieces of tissue placed within the coronary artery samples were in fact vein tissue. Supplementary figure 2 displays the results of a tSNE dimension reduction plot from which we identified two subgroups of interest for further review. One cluster displays abnormally high levels of expression of the ADIPOQ gene relative to the other subjects in these data. Consistent to our prior observation of poor correlation of heterogenous elements between histology and gene expression in GTEx, this ADIPOQ level did not correlate with the extent of adipose on the imaged sample (Ref! McCall AJHG). A review of the matched histology for the other highlighted subgroup of samples demonstrated that most exhibited features of vein tissue (thin-walled vessels with generalized collapse and no atherosclerosis) in at least one tissue segment in the image. In further study we sought to identify meaningful markers of vein tissue to identify whether the sequencing data for these samples comes from the artery or vein tissue in the histology, but no such markers were found. Without being able to be certain about which piece was selected for sequencing, as with samples that show one artery and one vein segment in the images, we chose to remove any potential vein samples from the analysis.

**Supplementary Figure 2.**
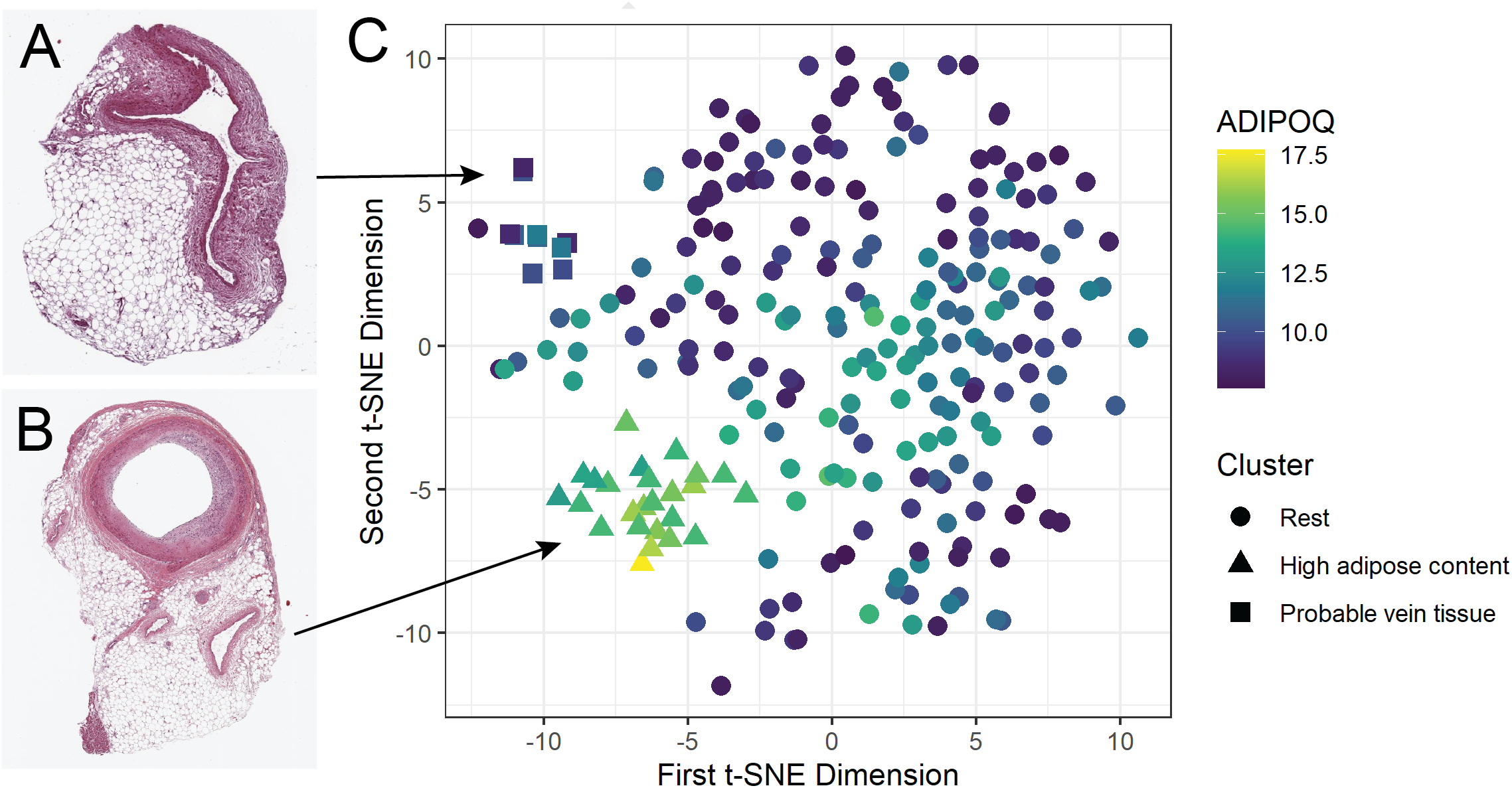
Dimension reduction on GTEx data suggests subgroups of samples with high adipose tissue composition and samples which are vein rather than artery.

### Supplementary Note 3: GTEx coronary artery histopathology

**Supplementary Figure 3.**
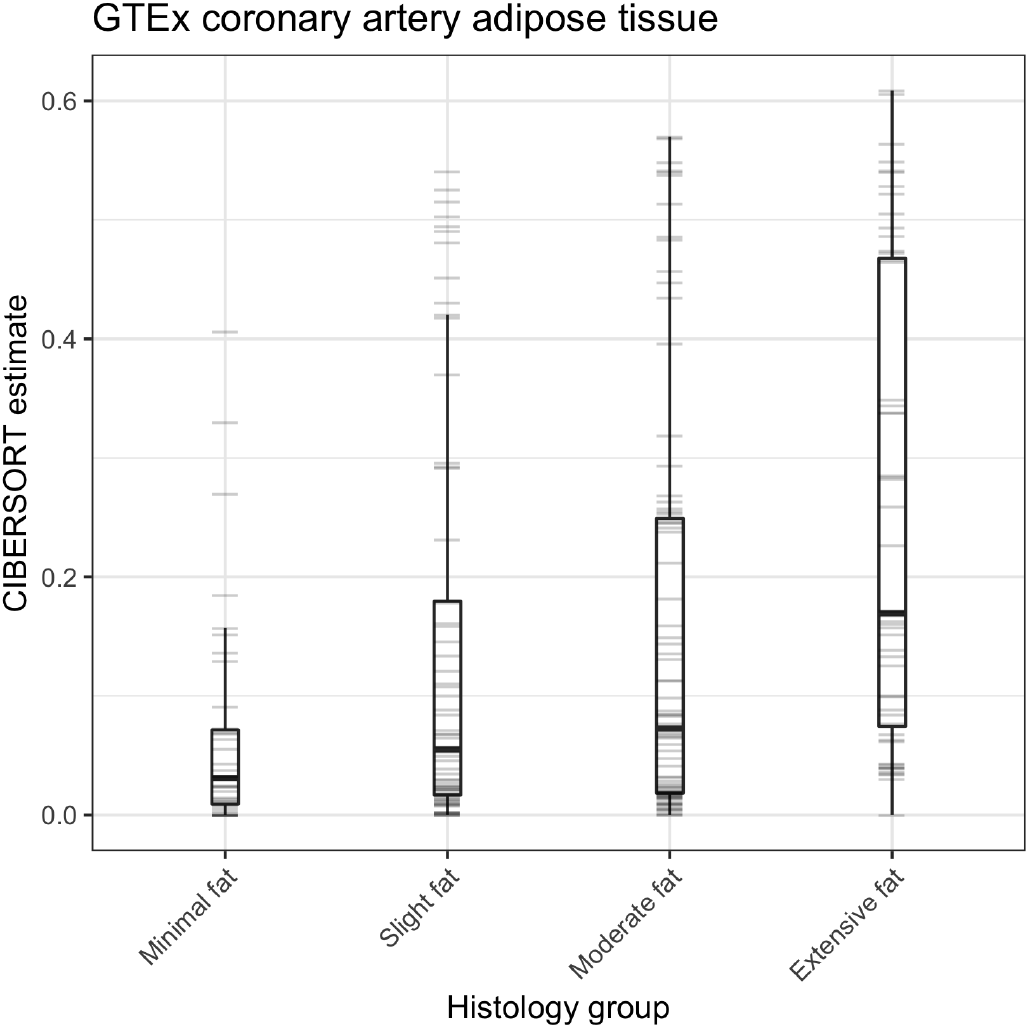
Distribution of CIBERSORT adipose estimates within histology groups.

**Supplementary Figure 4.**
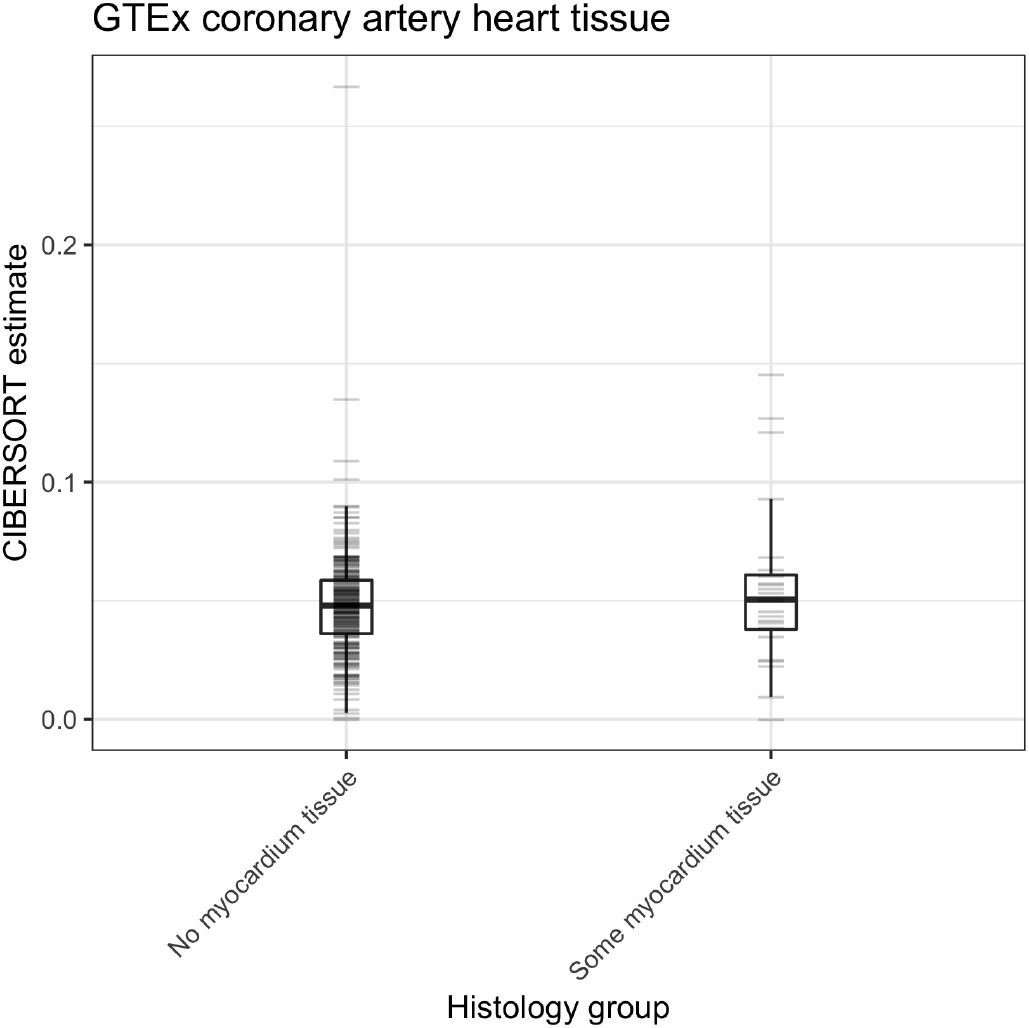
Distribution of CIBERSORT myocardium estimates within histology groups.

### Supplementary Note 4: CIBERSORT composition estimates neglect red blood cells

A notable omission from the CIBERSORT composition estimates is seen in the red blood cell type. These results do not reflect the expression of the genes specific to red blood cells, such as HBA1, HBA2, and HBB, in the GTEx data. These genes are entirely specific to red blood cells, so their expression signifies that they are in fact present in the tissues. Supplementary Fig. 5 shows the overall distribution of the VST expression counts of these genes in the GTEx tissues. We can clearly see that these genes are not only expressed in each of the tissue samples, but that there is a substantial degree of variance in these counts that should be reflected in the estimates as well.

**Supplementary Figure 5.**
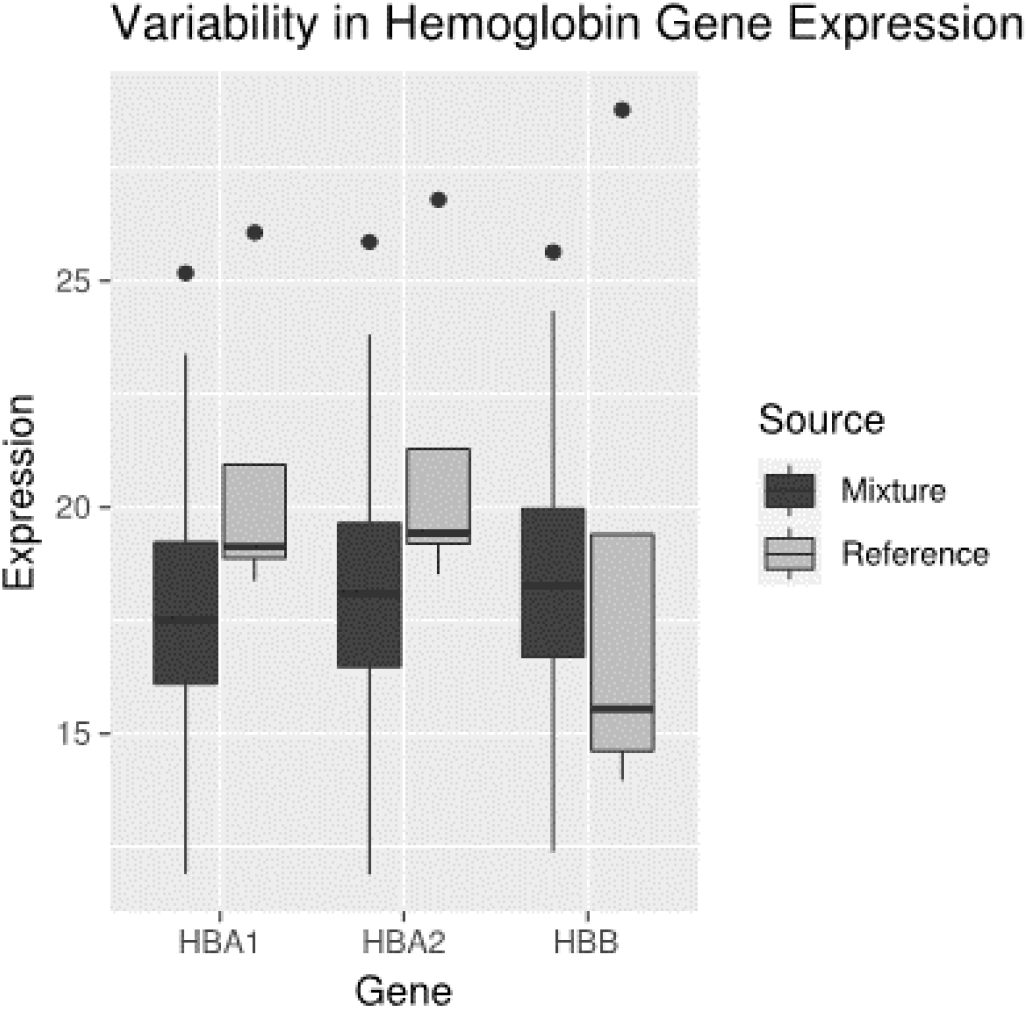
Distribution of hemoglobin genes in SRA and GTEx data.

### Supplementary Note 5: Distribution of effect sizes among DESeq2 results

**Supplementary Figure 6.**
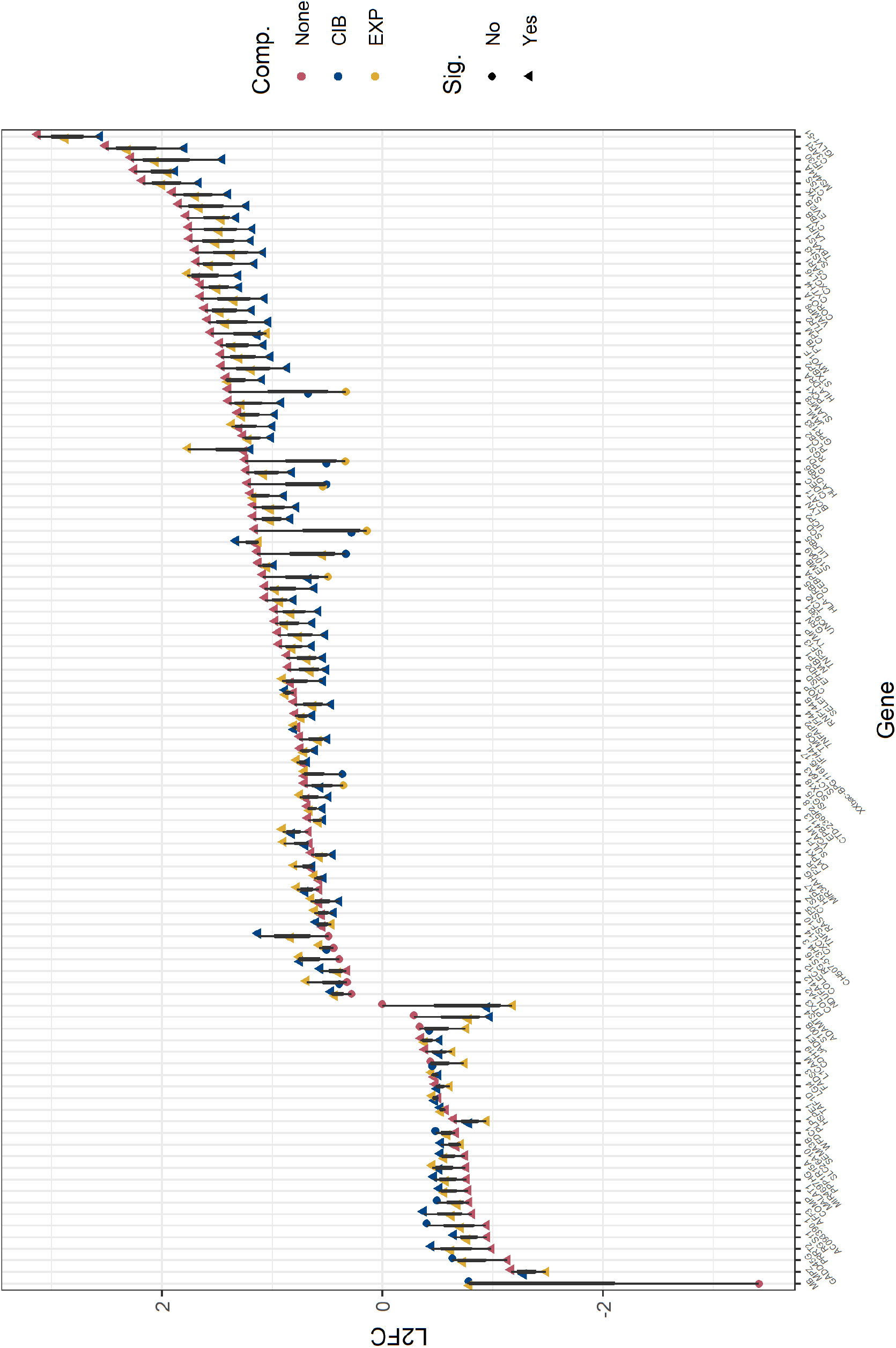
Distribution of effect sizes among transcripts found to be significant in at least one of the DESeq2 models, ordered to be increasing in the model that does not account for composition.

## Bibliography

1. Adeline R. Whitney, Maximilian Diehn, Stephen J. Popper, Ash A. Alizadeh, Jennifer C. Boldrick, David A. Relman, and Patrick O. Brown. Individuality and variation in gene expression patterns in human blood. Proceedings of the National Academy of Sciences of the United States of America, 100(4):1896–1901, 2003. ISSN 00278424. doi: 10.1073/pnas.252784499.

2. J. Perren Cobb, Michael N. Mindrinos, Carol Miller-Graziano, Steve E. Calvano, Henry V. Baker, Wenzhong Xiao, Krzysztof Laudanski, Bernard H. Brownstein, Constance M. Elson, Douglas L. Hayden, David N. Herndon, Stephen F. Lowry, Ronald V. Maier, David A. Schoenfeld, Lyle L. Moldawer, Ronald W. Davis, Ronald G. Tompkins, Paul Bankey, Timothy Billiar, David Camp, Irshad Chaudry, Bradley Freeman, Richard Gamelli, Nicole Gibran, Brian Harbrecht, Wyrta Heagy, David Heimbach, Jureta Horton, John Hunt, James Lederer, John Mannick, Bruce McKinley, Joseph Minei, Ernest Moore, Frederick Moore, Robert Munford, Avery Nathens, Grant O’keefe, Gary Purdue, Laurence Rahme, Daniel Remick, Matthew Sailors, Michael Shapiro, Geoffrey Silver, Richard Smith, Gregory Stephanopoulos, Gary Stormo, Mehmet Toner, Shaw Warren, Michael West, Steven Wolfe, and Vernon Young. Application of genome-wide expression analysis to human health and disease. Proceedings of the National Academy of Sciences of the United States of America, 102(13): 4801–4806, mar 2005. ISSN 00278424. doi: 10.1073/pnas.0409768102.

3. Chana Palmer, Maximilian Diehn, Ash A. Alizadeh, and Patrick O. Brown. Cell-type specific gene expression profiles of leukocytes in human peripheral blood. BMC Genomics, 7(1): 115, may 2006. ISSN 14712164. doi: 10.1186/1471-2164-7-115.

4. Liming Liang and William O.C. Cookson. Grasping nettles: Cellular heterogeneity and other confounders in epigenome-wide association studies. Human Molecular Genetics, 23(R1): 83–88, 2014. ISSN 14602083. doi: 10.1093/hmg/ddu284.

5. Andrew E. Jaffe and Rafael A. Irizarry. Accounting for cellular heterogeneity is critical in epigenome-wide association studies. Genome Biology, 15(2), feb 2014. ISSN 1474760X. doi: 10.1186/gb-2014-15-2-r31.

6. Baqer A. Haider, Alexander S. Baras, Matthew N. McCall, Joshua A. Hertel, Toby C. Cornish, and Marc K. Halushka. A critical evaluation of microRNA biomarkers in non-neoplastic disease. PLoS ONE, 9(2), feb 2014. ISSN 19326203. doi: 10.1371/journal.pone.0089565.

7. Oliver A. Kent, Matthew N. McCall, Toby C. Cornish, and Marc K. Halushka. Lessons from miR-143/145: The importance of cell-type localization of miRNAs, jul 2014. ISSN 13624962.

8. Yun Zhang, Jonavelle Cuerdo, Marc K Halushka, and Matthew N McCall. The effect of tissue composition on gene co-expression. Briefings in Bioinformatics, 00(January):1–13, 2019. ISSN 1467-5463. doi: 10.1093/bib/bbz135.

9. Aaron M. Newman, Chih Long Liu, Michael R. Green, Andrew J. Gentles, Weiguo Feng, Yue Xu, Chuong D. Hoang, Maximilian Diehn, and Ash A. Alizadeh. Robust enumeration of cell subsets from tissue expression profiles. Nature Methods, 12(5):453–457, 2015. ISSN 15487105. doi: 10.1038/nmeth.3337.

10. Ziyi Li and Hao Wu. Toast: improving reference-free cell composition estimation by cross-cell type differential analysis. Genome Biology, 20(1):18, 2019.

11. Ziyi Li, Zhijin Wu, Peng Jin, and Hao Wu. Dissecting differential signals in high-throughput data from complex tissues. Bioinformatics, 35(20):7803, 2019.

12. Xuran Wang, Jihwan Park, Katalin Susztak, Nancy R. Zhang, and Mingyao Li. Bulk tissue cell type deconvolution with multi-subject single-cell expression reference. Nature Communications, 10(1):1–9, dec 2019. ISSN 20411723. doi: 10.1038/s41467-018-08023-x.

13. GTEx Consortium. The genotype-tissue expression (gtex) project. Nature genetics, 45: 580–5, 6 2013. ISSN 1546-1718. doi: 10.1038/ng.2653.

14. GTEx Consortium. The gtex consortium atlas of genetic regulatory effects across human tissues. Science (New York, N.Y.), 369:1318–1330, 2020. ISSN 1095-9203. doi: 10.1126/science.aaz1776.

15. Joseph D Barry, Maud Fagny, Joseph N Paulson, Hugo J W L Aerts, John Platig, and John Quackenbush. Histopathological image qtl discovery of immune infiltration variants. iScience, 5:80–89, 7 2018. ISSN 2589-0042. doi: 10.1016/j.isci.2018.07.001.

16. Paul Gallins, Ehsan Saghapour, and Yi-Hui Zhou. Exploring the limits of combined image/’omics analysis for non-cancer histological phenotypes. Frontiers in genetics, 11: 555886, 2020. ISSN 1664-8021. doi: 10.3389/fgene.2020.555886.

17. Tim O Nieuwenhuis, Stephanie Y Yang, Rohan X Verma, Vamsee Pillalamarri, Dan E Arking, Avi Z Rosenberg, Matthew N McCall, and Marc K Halushka. Consistent rna sequencing contamination in gtex and other data sets. Nature communications, 11:1933, 2020. ISSN 2041-1723. doi: 10.1038/s41467-020-15821-9.

18. Matthew N McCall, Peter B Illei, and Marc K Halushka. Complex sources of variation in tissue expression data: Analysis of the gtex lung transcriptome. American journal of human genetics, 99:624–635, 2016. ISSN 1537-6605. doi: 10.1016/j.ajhg.2016.07.007.

19. Gen Li, Dereje Jima, Fred A Wright, and Andrew B Nobel. Ht-eqtl: integrative expression quantitative trait loci analysis in a large number of human tissues. BMC bioinformatics, 19: 95, 2018. ISSN 1471-2105. doi: 10.1186/s12859-018-2088-3.

20. Camila M Lopes-Ramos, Joseph N Paulson, Cho-Yi Chen, Marieke L Kuijjer, Maud Fagny, John Platig, Abhijeet R Sonawane, Dawn L DeMeo, John Quackenbush, and Kimberly Glass. Regulatory network changes between cell lines and their tissues of origin. BMC genomics, 18:723, 9 2017. ISSN 1471-2164. doi: 10.1186/s12864-017-4111-x.

21. Nicole M Ferraro, Benjamin J Strober, Jonah Einson, Nathan S Abell, Francois Aguet, Alvaro N Barbeira, Margot Brandt, Maja Bucan, Stephane E Castel, Joe R Davis, Emily Greenwald, Gaelen T Hess, Austin T Hilliard, Rachel L Kember, Bence Kotis, YoSon Park, Gina Peloso, Shweta Ramdas, Alexandra J Scott, Craig Smail, Emily K Tsang, Seyedeh M Zekavat, Marcello Ziosi, Aradhana, TOPMed Lipids Working Group, Kristin G Ardlie, Themistocles L Assimes, Michael C Bassik, Christopher D Brown, Adolfo Correa, Ira Hall, Hae Kyung Im, Xin Li, Pradeep Natarajan, GTEx Consortium, Tuuli Lappalainen, Pejman Mohammadi, Stephen B Montgomery, and Alexis Battle. Transcriptomic signatures across human tissues identify functional rare genetic variation. Science (New York, N.Y.), 369, 2020. ISSN 1095-9203. doi: 10.1126/science.aaz5900.

22. Renu Virmani, Frank D. Kolodgie, Allen P. Burke, Andrew Farb, and Stephen M. Schwartz. Lessons from sudden coronary death: A comprehensive morphological classification scheme for atherosclerotic lesions. Arteriosclerosis, Thrombosis, and Vascular Biology, 20(5):1262–1275, 2000. ISSN 10795642. doi: 10.1161/01.ATV.20.5.1262.

23. Christopher Wilks, Leonardo Collado-Torres, Shijie C. Zheng, Andrew E. Jaffe, Abhinav Nellore, Kasper D. Hansen, and Ben Langmead. recount3 pre-print title todo. bioRxiv, 2020. doi: 10.1101/TODO.

24. Leonardo Collado-Torres. Explore and download data from the recount3 project, 2021. https://github.com/LieberInstitute/recount 3 - R package version 1.0.7.

25. Matthew N Bernstein, AnHai Doan, and Colin N Dewey. MetaSRA: normalized human sample-specific metadata for the Sequence Read Archive. Bioinformatics, 33(18):2914–2923, sep 2017. ISSN 1367-4803. doi: 10.1093/bioinformatics/btx334.

26. Davide Risso, John Ngai, Terence P. Speed, and Sandrine Dudoit. Normalization of RNA-seq data using factor analysis of control genes or samples. Nature Biotechnology, 32(9): 896–902, 2014. ISSN 15461696. doi: 10.1038/nbt.2931.

27. Hao Wu, Chi Wang, and Zhijin Wu. PROPER: comprehensive power evaluation for differential expression using RNA-seq. Bioinformatics, 31(2):233–241, jan 2015. ISSN 1460-2059. doi: 10.1093/bioinformatics/btu640.

28. Michael I. Love, Wolfgang Huber, and Simon Anders. Moderated estimation of fold change and dispersion for RNA-seq data with DESeq2. Genome Biology, 15(12):1–21, 2014. ISSN 1474760X. doi: 10.1186/s13059-014-0550-8.

29. L.J.P. van der Maaten and G.E. Hinton. Visualizing high-dimensional data using t-sne. Journal of Machine Learning Research, 9:2579–2605, 2008.

30. L.J.P. van der Maaten. Accelerating t-sne using tree-based algorithms. Journal of Machine Learning Research, 15:3221–3245, 2014.

31. Jesse H. Krijthe. Rtsne: T-Distributed Stochastic Neighbor Embedding using Barnes-Hut Implementation, 2015. R package version 0.15.

32. Marie A.C. Depuydt, Koen H.M. Prange, Lotte Slenders, Tiit Örd, Danny Elbersen, Arjan Boltjes, Saskia C.A. De Jager, Folkert W. Asselbergs, Gert J. De Borst, Einari Aavik, Tapio Lönnberg, Esther Lutgens, Christopher K. Glass, Hester M. Den Ruijter, Minna U. Kaikkonen, Ilze Bot, Bram Slütter, Sander W. Van Der Laan, Seppo Yla-Herttuala, Michal Mokry, Johan Kuiper, Menno P.J. De Winther, and Gerard Pasterkamp. Microanatomy of the Human Atherosclerotic Plaque by Single-Cell Transcriptomics. Circulation Research, 127: 1437–1455, 2020. ISSN 15244571. doi: 10.1161/CIRCRESAHA.120.316770.

33. Colin Megill, Bruce Martin, Charlotte Weaver, Sidney Bell, Lia Prins, Seve Badajoz, Brian McCandless, Angela Oliveira Pisco, Marcus Kinsella, Fiona Griffin, Justin Kiggins, Genevieve Haliburton, Arathi Mani, Matthew Weiden, Madison Dunitz, Maximilian Lombardo, Timmy Huang, Trent Smith, Signe Chambers, Jeremy Freeman, Jonah Cool, and Ambrose Carr. cellxgene: a performant, scalable exploration platform for high dimensional sparse matrices. bioRxiv, 2021. doi: 10.1101/2021.04.05.438318.

34. Anna C. Neufeld, Lucy L. Gao, and Daniela M. Witten. Tree-values: selective inference for regression trees. 6 2021.

