## Supplementary material for "Considerations for Deconvolution: A Case Study with GTEx Coronary Artery Tissues": Data and R Code: workflow.Rmd

````
---
title: "Atherosclerosis Deconvolution Paper Code"
author: "Zach Brehm"
date: "5/17/2022"
output: html_document
---

```{r setup, include=FALSE}
knitr::opts_chunk$set(echo = TRUE)
library(DESeq2)
library(tidyverse)
library(ggmnmlab)
```

## Getting the GTEx data

We used the recount3 package to download the GTEx coronary artery data. These data are
available as a part of the overall blood vessel set found in recount3.

```{r download gtex data}
library(recount3)
human_projects <- available_projects()
proj_info <- subset(
  human_projects,
  project == "BLOOD_VESSEL" & project_type == "data_sources"
)
rse_coronary <- create_rse(proj_info)
rse_coronary <- rse_coronary[, rse_coronary$gtex.smtsd == "Artery - Coronary"]
assay(rse_coronary, "counts") <- transform_counts(rse_coronary)
saveRDS(rse_coronary, file = "data/rse_coronary_recount3_full.Rds")
```

There are several subjects with duplicated samples in the coronary artery data so we
subset these data to only include one sample per subject. Features with counts less than
1,000 in all subjects are removed as well. Numerous sample qualities are added to the column data
to account for adjacent histology and demographic information in the GTEx samples.

```{r filter low counts and duplicate subjects}
assays(rse_coronary)[1] <- NULL
lowCounts <- apply(assay(rse, "counts"), 1, function(x) {
  all(x < 1000)
})

rse_coronary <- rse_coronary[-which(lowCounts), ]

gtexNames <- str_split(cibAbs$InputSample, "-") %>%
  sapply(FUN = function(x) paste(x[1], x[2], x[3], sep = "-"))
cibAbs$InputSample <- gtexNames
rse_coronary$histologyID <- gtexNames

duplicatedSamples <- which(duplicated(rse_coronary$gtex.subjid))

## duplicated samples results 88, 120, 212, 213, 214
## 49 GTEX-P4PP-3026 (same subject for 88 and 120, both vein in histology)
## 22 GTEX-QV44-0726 (same subject for 212 which has NA SRR)
## 96 GTEX-QEG5-1026 (same subject for 213 which has NA SRR)
## 32 GTEX-QLQ7-0326 (same subject for 214 which has NA SRR)

rse_coronary <- rse_coronary[, -duplicatedSamples]

cac <- read_excel("Coronary_artery_case_images_8.27.20.xlsx")

cacSub <- subset(cac, `Specimen ID` %in% rse_coronary$histologyID)
## one duplicated Specimen ID
cacSub <- cacSub[-148, ]

map <- match(rse_coronary$histologyID, cacSub$`Specimen ID`)

cacSub <- cacSub[map, ]

## confirm that these tables are aligned, following line should evalute as TRUE

identical(rse_coronary$histologyID, cacSub$`Specimen ID`)

rse_coronary$virmani <- cacSub$`Plaque Type`

rse_coronary$plaque <- factor(rse_coronary$virmani)

rse_coronary$plaque[!(rse_coronary$virmani == 0)] <- 1

rse_coronary$virmani <- factor(rse_coronary$virmani)

rse_coronary$plaque <- factor(rse_coronary$plaque)

rse_coronary$year <- parse_date_time(rse_coronary$gtex.smgebtchd, "mdy") %>%
  year() %>%
  factor()

rse_coronary$age <- factor(rse_coronary$gtex.age, labels = 0:5)

rse_coronary$sex <- factor(rse_coronary$gtex.sex, labels = c("Male", "Female"))

veinSamples <- c(
  "GTEX-15DDE-1226",
  "GTEX-P4PP-3026",
  "GTEX-132AR-1326",
  "GTEX-15TU5-1226",
  "GTEX-RTLS-0726",
  "GTEX-12ZZX-0626",
  "GTEX-1CB4E-0726",
  "GTEX-15CHR-0526",
  "GTEX-ZYT6-0826",
  "GTEX-12WSC-0926",
  "GTEX-14BMV-0926",
  "GTEX-17GQL-0426",
  "GTEX-ZDYS-0126"
)

veinInd <- which(rse_coronary$histologyID %in% veinSamples)

rse_coronary <- rse_coronary[, -veinInd]

saveRDS(rse_coronary,
  file = "data/rse_coronary_recount3_for_cibersort.Rds"
)
```

## Getting the reference data

```{r}
library(recount3)

# load in the samples of interest, their respective celltypes,
# and the project IDs that they are found in.

projects <- read.table("data/vals_srp.txt")
runs <- read.table("data/vals_srr.txt")
celltypes <- read.table("data/celltypes.txt")

# identify the projects available in recount3.
human_projects <- available_projects()

projects <- projects[projects %in% human_projects$project]

project <- subset(
  human_projects,
  project == projects[1] & project_type == "data_sources"
)

rse <- create_rse(project_info = project)

projects <- projects[-1]

for (i in 1:length(projects)) {
  project <- subset(human_projects, project == projects[i] & project_type == "data_sources")
  tmp_rse <- create_rse(project_info = project)
  rse <- cbind(rse, tmp_rse)
}

rse <- rse[, colnames(rse) %in% runs]

map <- match(runs, colnames(rse))

rse <- rse[, map]

rse$celltype <- celltypes

assay(rse, "counts") <- transform_counts(rse)

saveRDS(rse, file = "data/rse_reference_full.Rds")

duplicated_srs <- unique(rse$sra.sample_acc.x[duplicated(rse$sra.sample_acc.x)])

dup_srs_ind <- sapply(duplicated_srs, function(x) which(rse$sra.sample_acc.x == x))

counts_mat <- assay(rse, "counts")

rep_sums <- lapply(dup_srs_ind, function(x) rowSums(counts_mat[, x]))

dup_remove <- sapply(1:length(dup_srs_ind), function(x) dup_srs_ind[[x]][-1]) %>% unlist()

dup_replace <- sapply(1:length(dup_srs_ind), function(x) dup_srs_ind[[x]][1])

for (i in 1:length(rep_sums)) {
  counts_mat[, dup_replace[i]] <- rep_sums[[i]]
}

counts_mat <- counts_mat[, -dup_remove]

rse <- rse[, -dup_remove]

assay(rse, "counts") <- counts_mat

# raw counts not necessary for analyses, remove this assay to conserve memory

assays(rse)[1] <- NULL

lowCounts <- apply(assay(rse), 1, function(x) {
  all(x < 1000)
})

rse <- rse[-which(lowCounts), ]

saveRDS(rse, file = "data/rse_reference_no_reps.Rds")

rse <- rse[, -which(rse$celltype == "Erythrocyte")]

saveRDS(rse, file = "data/rse_reference_noRBC.Rds")
```

## Exploratory analysis GTEx

```{r}
gtex <- readRDS(file = "data/rse_coronary_recount3.Rds")

cac <- read_excel("data/Coronary_artery_case_images_8.27.20.xlsx")

gtexNames <- str_split(colnames(gtex), "-") %>% sapply(FUN = function(x) paste(x[1], x[2], x[3], sep = "-"))

cacSub <- subset(cac, `Specimen ID` %in% gtexNames)
cacSub <- cacSub[-148, ]

map <- match(gtexNames, cacSub$`Specimen ID`)

cacSub <- cacSub[map, ]

lowCounts <- apply(assay(gtex), 1, function(x) {
  all(x < 1000)
})
gtex <- gtex[-which(lowCounts), ]

gtex$Plaque <- factor(cacSub$`Plaque Type`)
gtex$Inflammation <- factor(cacSub$Inflammation)
gtex$Fat <- factor(cacSub$Fat)
gtex$Myocardium <- factor(cacSub$Myocardium)
gtex$PlaqueInd <- factor((cacSub$`Plaque Type` > 0) * 1)

gtexDds <- DESeqDataSet(gtex, design = ~1)
gtexVst <- varianceStabilizingTransformation(gtexDds)
gtexDist <- dist(t(assay(gtexVst)))

veinSamples <- c(
  "GTEX-15DDE-1226",
  "GTEX-P4PP-3026",
  "GTEX-132AR-1326",
  "GTEX-15TU5-1226",
  "GTEX-RTLS-0726",
  "GTEX-12ZZX-0626",
  "GTEX-1CB4E-0726",
  "GTEX-15CHR-0526",
  "GTEX-ZYT6-0826",
  "GTEX-12WSC-0926",
  "GTEX-14BMV-0926",
  "GTEX-17GQL-0426",
  "GTEX-ZDYS-0126"
)

namesC1 <- c(
  "GTEX-YB5K-0426",
  "GTEX-17GQL-0426",
  "GTEX-ZPIC-0326",
  "GTEX-ZDYS-0126",
  "GTEX-13W3W-0626",
  "GTEX-1IKOE-0826",
  "GTEX-13QBU-0126",
  "GTEX-YF7O-0226",
  "GTEX-12WSE-1126",
  "GTEX-1BAJH-1226",
  "GTEX-1AYCT-0626",
  "GTEX-WEY5-0326",
  "GTEX-11I78-1026",
  "GTEX-14PN3-0426",
  "GTEX-ZG7Y-2326",
  "GTEX-139YR-0726",
  "GTEX-14E1K-0926",
  "GTEX-1F5PK-0426",
  "GTEX-1HFI7-0626",
  "GTEX-1GL5R-0226",
  "GTEX-YEC3-0326",
  "GTEX-ZZPU-1026",
  "GTEX-1B97J-0826",
  "GTEX-ZQG8-0626"
)

veinID <- which(gtexNames %in% veinSamples)
clusterLabs <- factor(
  x = rep("Remaining", length(gtexNames)),
  levels = c("Remaining", "High adipose content", "Probable vein tissue")
)
clusterLabs[which(gtexNames %in% namesC1)] <- "High adipose content"
clusterLabs[which(gtexNames %in% veinSamples)] <- "Probable vein tissue"

#set.seed(z)

gtexTsneDist <- Rtsne(gtexDist,
  perplexity = 40, theta = 0,
  pca = FALSE, is_distance = TRUE
)

tsneDistDf <- data.frame(
  X = gtexTsneDist$Y[, 1],
  Y = gtexTsneDist$Y[, 2],
  Plaque = gtexVst$Plaque,
  PlaqueInd = gtexVst$PlaqueInd,
  Sample = gtexNames,
  Inflammation = gtexVst$Inflammation,
  Sex = factor(gtexVst$gtex.sex, labels = c("Male", "Female")),
  Age = factor(gtexVst$gtex.age),
  Autolysis = factor(gtexVst$gtex.dthhrdy),
  Fat = gtexVst$Fat,
  Myocardium = gtexVst$Myocardium,
  ADIPOQ = assay(gtexVst)[which(rowData(gtexVst)$gene_name == "ADIPOQ"), ],
  TTN = assay(gtexVst)[which(rowData(gtexVst)$gene_name == "TTN"), ],
  Cluster = clusterLabs
)

ggtsnedist <- ggplot(tsneDistDf, aes(x = X, y = Y, color = ADIPOQ, text = Sample, shape = Cluster)) +
  geom_point(size = 3) +
  labs( # title = "Dimension Reduction with Rtsne",
    # subtitle = "Subpopulations of interest",
    x = "First t-SNE Dimension",
    y = "Second t-SNE Dimension",
    color = "ADIPOQ"
  ) +
  guides(alpha = "none") +
  scale_color_viridis_c() + ## with end = 1 the yellow is a bit too bright for a white background
  theme_bw()
ggtsnedist
ggsave(
  filename = "figures/tsne_vein_clusters.png", plot = ggtsnedist,
  device = "png", dpi = "print", width = 6, height = 4, units = "in"
)

gtex_vst <- DESeqDataSet(gtex, design = ~1) %>% varianceStabilizingTransformation(.)

gtex_dds <- DESeqDataSet(gtex, design = ~plaque) %>% DESeq()

plaque_results <- results(gtex_dds, contrast = c("plaque", 1, 0))

plaque_results$HGNC <- rowData(gtex_dds)$gene_name

plaque_results <- plaque_results[-which(plaque_results$baseMean < 500), ]

plaque_results <- plaque_results[order(abs(plaque_results$log2FoldChange), decreasing = TRUE), ]

sort_abs_lfc <- plaque_results$HGNC[order(abs(plaque_results$log2FoldChange), decreasing = TRUE)]

pcaGtex <- prep(t(assay(gtex_vst)), center = TRUE, scale = "none") %>%
  pca(method = "svd", nPcs = 20)

hgncVec <- rowData(gtex_dds)$gene_name

pcaGtex_top100 <- prep(t(assay(gtex_vst)[which(rowData(gtex_vst)$gene_name %in% sort_abs_lfc[1:100]), ]),
  center = TRUE, scale = "none"
) %>% pca(method = "svd", nPcs = 20)

scores <- data.frame(pcaGtex@scores) %>%
  mutate(
    `Plaque Score` = gtex_vst$virmani,
    `Macrophage Score` = gtex_vst$MacScore,
    `Plaque` = gtex_vst$plaque,
    `Macrophage` = gtex_vst$MacInd,
    Sex = gtex_vst$sex,
    Name = rownames(data.frame(pcaGtex@scores))
  )

scores_top100 <- data.frame(pcaGtex_top100@scores) %>%
  mutate(
    `Plaque Score` = gtex_vst$virmani,
    `Macrophage Score` = gtex_vst$MacScore,
    `Plaque` = gtex_vst$plaque,
    `Macrophage` = gtex_vst$MacInd,
    Sex = gtex_vst$sex,
    Name = rownames(data.frame(pcaGtex_top100@scores))
  )

gg_pca_gtex <- ggplot(scores, aes(x = PC1, y = PC2, color = Plaque)) +
  geom_point() +
  scale_color_mnmlab(palette = "highContrast") +
  labs(
    title = "PCA of GTEx coronary artery",
    subtitle = "Full set of 235 samples"
  ) +
  theme_bw()
ggsave(
  filename = "figures/gg_pca_gtex.png", plot = gg_pca_gtex,
  device = "png", dpi = "print", width = 5, height = 5, units = "in"
)

gg_pca_gtex_top100 <- ggplot(scores_top100, aes(x = PC1, y = PC2, color = Sex)) +
  geom_point() +
  scale_color_mnmlab(palette = "highContrast") +
  labs(
    # title = "PCA of GTEx coronary artery",
    # subtitle = "Full set of 235 samples"
  ) +
  theme_bw()
ggsave(
  filename = "figures/gg_pca_gtex_top100.png", plot = gg_pca_gtex_top100,
  device = "png", dpi = "print", width = 5, height = 5, units = "in"
)

markerGenes <- readRDS("data/markerGenes.Rds")

smcInd <- which(hgncVec %in% markerGenes$SMC)

adiInd <- which(hgncVec %in% markerGenes$ADI)

rbcInd <- which(hgncVec %in% markerGenes$RBC)

cmcInd <- which(hgncVec %in% cmcGenes)

endInd <- which(hgncVec %in% endGenes)

lymInd <- which(hgncVec %in% markerGenes$LYM)

macInd <- which(hgncVec %in% markerGenes$MAC)

ighInd <- c(
  which(startsWith(hgncVec, "IGHV")),
  which(startsWith(hgncVec, "IGHJ")),
  which(startsWith(hgncVec, "IGHD")),
  which(startsWith(hgncVec, "IGHE")),
  which(startsWith(hgncVec, "IGHM")),
  which(startsWith(hgncVec, "IGHA")),
  which(startsWith(hgncVec, "IGHG"))
)

igkInd <- c(
  which(startsWith(hgncVec, "IGKV")),
  which(startsWith(hgncVec, "IGKJ")),
  which(startsWith(hgncVec, "IGKC"))
)

iglInd <- c(
  which(startsWith(hgncVec, "IGLV")),
  which(startsWith(hgncVec, "IGLJ")),
  which(startsWith(hgncVec, "IGLC"))
)

igInd <- c(ighInd, igkInd, iglInd)

select <- c(macInd, lymInd, smcInd, endInd, adiInd, cmcInd, rbcInd)

gtexSub <- DESeqDataSet(gtexSub, design = ~1) %>%
  estimateSizeFactors() %>%
  estimateDispersions()
gtexCounts <- counts(gtexSub, normalized = TRUE)
gtexCounts <- gtexCounts[select, ]
rowMeanGtex <- rowMeans(gtexCounts)
rowSdsGtex <- rowSds(gtexCounts)
for (i in 1:nrow(gtexCounts)) {
  gtexCounts[i, ] <- (gtexCounts[i, ] - rowMeanGtex[i]) / rowSdsGtex[i]
}
rownames(gtexCounts) <- rowData(gtexSub)$gene_name[select]
celltypenames <- vector(length = nrow(gtexCounts))
celltypenames[1:6] <- "MAC"
celltypenames[7:29] <- "LYM"
celltypenames[30:37] <- "SMC"
celltypenames[38:41] <- "END"
celltypenames[42:44] <- "ADI"
celltypenames[45:55] <- "CMC"
celltypenames[56:58] <- "RBC"

celltypenames <- factor(celltypenames,
  levels = c(
    "MAC",
    "LYM",
    "SMC",
    "END",
    "ADI",
    "CMC",
    "RBC"
  )
)

haPlq <- HeatmapAnnotation(
  Plaque = factor(gtexSub$Plaque),
  col = list(
    "Plaque" = c(
      "0" = "#440154FF",
      "1" = "#472D7BFF",
      "2" = "#3B528BFF",
      "3" = "#2C728EFF",
      "4" = "#21908CFF",
      "5" = "#27AD81FF",
      "6" = "#5DC863FF",
      "7" = "#AADC32FF",
      "8" = "#FDE725FF"
    )
  ),
  annotation_name_side = "right"
)

haMac <- HeatmapAnnotation(
  `Macrophage Score` = factor(gtexSub$MacInd),
  col = list(
    `Macrophage Score` = c(
      "0" = "#0077BB",
      "1" = "#33BBEE",
      "2" = "#009988",
      "3" = "#EE7733",
      "4" = "#CC3311",
      "5" = "#EE3377"
    )
  ),
  annotation_name_side = "right"
)

htPlq <- Heatmap(gtexCounts,
  column_order = order(gtexSub$Plaque),
  column_split = gtexSub$MacInd,
  row_split = celltypenames,
  cluster_row_slices = FALSE,
  column_names_gp = gpar(fontsize = 7),
  row_names_gp = gpar(fontsize = 5),
  top_annotation = haBot,
  bottom_annotation = ha,
  name = "Gene expression z-score",
  cluster_rows = FALSE,
  cluster_columns = FALSE,
  column_labels = rep("", 235),
  heatmap_legend_param = list(
    nrow = 3,
    ncol = 1
  )
)
draw(ht1)

htMac <- Heatmap(gtexCounts,
  column_order = order(gtexSub$MacInd),
  column_split = gtexSub$Plaque,
  row_split = celltypenames,
  cluster_row_slices = FALSE,
  column_names_gp = gpar(fontsize = 7),
  row_names_gp = gpar(fontsize = 5),
  top_annotation = ha,
  bottom_annotation = haBot,
  name = "Gene expression z-score",
  cluster_rows = FALSE,
  cluster_columns = FALSE,
  column_labels = rep("", 236),
  heatmap_legend_param = list(
    nrow = 3,
    ncol = 1
  )
)
draw(ht2)

gtex$LymScore <- (colSums(assay(gtex_vst)[lymInd, ]) - mean(colSums(assay(gtex_vst)[lymInd, ]))) / sd(colSums(assay(gtex_vst)[lymInd, ]))
gtex$RbcScore <- (colSums(assay(gtex_vst)[rbcInd, ]) - mean(colSums(assay(gtex_vst)[rbcInd, ]))) / sd(colSums(assay(gtex_vst)[rbcInd, ]))
gtex$MacScore <- (colSums(assay(gtex_vst)[macInd, ]) - mean(colSums(assay(gtex_vst)[macInd, ]))) / sd(colSums(assay(gtex_vst)[macInd, ]))
gtex$CmcScore <- (colSums(assay(gtex_vst)[cmcInd, ]) - mean(colSums(assay(gtex_vst)[cmcInd, ]))) / sd(colSums(assay(gtex_vst)[cmcInd, ]))
gtex$SmcScore <- (colSums(assay(gtex_vst)[smcInd, ]) - mean(colSums(assay(gtex_vst)[smcInd, ]))) / sd(colSums(assay(gtex_vst)[smcInd, ]))
gtex$AdiScore <- (colSums(assay(gtex_vst)[adiInd, ]) - mean(colSums(assay(gtex_vst)[adiInd, ]))) / sd(colSums(assay(gtex_vst)[adiInd, ]))
gtex$EndScore <- (colSums(assay(gtex_vst)[endInd, ]) - mean(colSums(assay(gtex_vst)[endInd, ]))) / sd(colSums(assay(gtex_vst)[endInd, ]))

gtex$MacInd <- rep(0, ncol(gtex))
gtex$MacInd[which(gtex$MacScore > -1.75716507 & gtex$MacScore <= -0.64328696)] <- 1
gtex$MacInd[which(gtex$MacScore <= -0.04876403 & gtex$MacScore > -0.64328696)] <- 2
gtex$MacInd[which(gtex$MacScore > -0.04876403 & gtex$MacScore <= 0.59118248)] <- 3
gtex$MacInd[which(gtex$MacScore <= 2.16349136 & gtex$MacScore > 0.59118248)] <- 4
gtex$MacInd[which(gtex$MacScore > 2.16349136)] <- 5

subsetInd45 <- which(gtex$virmani %in% c(1:8) & gtex$MacScore > -0.04876403)
subsetInd01 <- which(gtex$virmani == 0 & gtex$MacScore <= -0.04876403)

gtexSub045 <- gtex[, c(subsetInd01, subsetInd45)]

gtex_vst045 <- DESeqDataSet(gtexSub045, design = ~1) %>% varianceStabilizingTransformation(.)

saveRDS(gtex, file = "data/gtex_n235_exp_scores.Rds")


pcaGtex045 <- prep(t(assay(gtex_vst045)), center = TRUE, scale = "none") %>%
  pca(method = "svd", nPcs = 20)

scores045 <- data.frame(pcaGtex045@scores) %>%
  mutate(
    `Plaque Score` = gtex_vst045$virmani,
    `Macrophage Score` = gtex_vst045$MacScore,
    `Plaque` = gtex_vst045$plaque,
    `Macrophage` = gtex_vst045$MacInd,
    Name = rownames(data.frame(pcaGtex045@scores))
  )

gg_pca_gtexSub <- ggplot(scores045, aes(x = PC1, y = PC2, color = Plaque)) +
  geom_point() +
  scale_color_mnmlab(palette = "highContrast") +
  labs(
    title = "PCA of GTEx coronary artery",
    subtitle = "Reduced set of 138 samples"
  ) +
  theme_bw()
ggsave(
  filename = "figures/gg_pca_gtexSub.png", plot = gg_pca_gtexSub,
  device = "png", dpi = "print", width = 5, height = 5, units = "in"
)
```

## Exploratory analysis SRA

```{r}
library(Rtsne)

## this script is to perform dimension reduction of gene expression data
## for plotting using the cmd method and tsne method

## read in batch corrected gene expression data
rse <- readRDS(file = "data/ruv_reference.Rds")

## make rse into dds object for variance stabilizing
dds <- DESeqDataSet(rse, design = ~1) %>% varianceStabilizingTransformation(.)

## create vst object to attenuate the mean/variance relationship
## of the expression data
# vst <- varianceStabilizingTransformation(dds)

## compute euclidean distance between samples
dist <- dist(t(assay(dds)))

pca_rse <- prep(t(assay(dds)), center = TRUE, scale = "pareto") %>%
  pca(method = "svd", nPcs = 50)

pcaDf <- data.frame(pca_rse@scores) %>% mutate(Celltype = cmdDf$Celltype)

gg_pca <- ggplot(pcaDf, aes(x = PC1, y = PC2, color = Celltype, alpha = 0.8)) +
  geom_point() +
  labs( # title = "Reference data for CIBERSORT",
    # subtitle = "Dimension reduction with cmdscale",
    x = "PC1",
    y = "PC2",
    color = "Celltype"
  ) +
  scale_color_mnmlab(palette = "martink8") +
  guides(alpha = "none") +
  theme_bw()

gg_pca

## compute multidimensional scaling of distance matrix
## and convert to dataframe to plot
cmdDf <- cmdscale(dist) %>%
  data.frame(.) %>%
  rownames_to_column(var = "Sample") %>%
  mutate(
    Celltype = dds$celltype,
    Project = dds$study
  ) %>%
  select(
    Sample,
    Celltype,
    Project,
    everything()
  )

cmdDf$Celltype <- factor(cmdDf$Celltype,
  levels = c(
    "FatCell",
    "CardiacMuscleCell",
    "Macrophage",
    "Lymphocytes",
    "EndothelialCell",
    "SmoothMuscleCell",
    "Erythrocyte"
  ),
  labels = c(
    "ADI",
    "CMC",
    "MAC",
    "LYM",
    "END",
    "SMC",
    "RBC"
  )
)

gg_cmd <- ggplot(cmdDf, aes(x = X1, y = X2, color = Celltype, alpha = 0.8, text = Project)) +
  geom_point() +
  labs(
    title = "Reference data for CIBERSORT",
    subtitle = "Dimension reduction with cmdscale",
    x = "First Dimension",
    y = "Second Dimension",
    color = "Celltype"
  ) +
  scale_color_mnmlab(palette = "martink8") +
  guides(alpha = "none") +
  theme_bw()

gg_cmd

ggsave(
  filename = "figures/gg_cmd.png", plot = gg_cmd,
  device = "png", dpi = "print", width = 5, height = 5, units = "in"
)

#set.seed(m)
## run tsne with the previously created distance matrix
cellTsne <- Rtsne(dist, is_distance = TRUE, perplexity = 40)

## make tsne dataframe and plot
tsneDf <- data.frame(
  X = cellTsne$Y[, 1],
  Y = cellTsne$Y[, 2],
  Z = cmdDf$Celltype,
  Project = dds$study
)

gg_tsne <- ggplot(tsneDf, aes(x = X, y = Y, color = Z, text = Project, alpha = 0.8)) +
  geom_point() +
  labs( # title = "Reference data for CIBERSORT",
    # subtitle = "Dimension reduction with Rtsne",
    x = "First t-SNE Dimension",
    y = "Second t-SNE Dimension",
    color = "Cell type",
    text = dds$study
  ) +
  guides(alpha = "none") +
  scale_color_mnmlab(
    palette = "martink8",
    labels = c(
      "ADI (n = 74)",
      "CMC (n = 33)",
      "MAC (n = 188)",
      "LYM (n = 380)",
      "END (n = 81)",
      "SMC (n = 31)",
      "RBC (n = 3)"
    )
  ) +
  theme_bw()

gg_tsne

ggsave(
  filename = "figures/gg_tsne.png", plot = gg_tsne,
  device = "png", dpi = "print", width = 5, height = 5, units = "in"
)

umap_rse <- umap(t(assay(dds)))

umapDf <- data.frame(
  X = umap_rse$layout[, 1],
  Y = umap_rse$layout[, 2],
  Z = cmdDf$Celltype
)


gg_umap <- ggplot(umapDf, aes(x = X, y = Y, color = Z, alpha = 0.8)) +
  geom_point() +
  labs( # title = "Reference data for CIBERSORT",
    # subtitle = "Dimension reduction with Rtsne",
    x = "First UMAP Dimension",
    y = "Second UMAP Dimension",
    color = "Cell type"
  ) +
  guides(alpha = "none") +
  scale_color_mnmlab(
    palette = "martink8",
    labels = c(
      "ADI (n = 74)",
      "CMC (n = 33)",
      "MAC (n = 188)",
      "LYM (n = 380)",
      "END (n = 81)",
      "SMC (n = 31)",
      "RBC (n = 3)"
    )
  ) +
  theme_bw()

gg_umap

ggsave(
  filename = "figures/gg_umap.png", plot = gg_umap,
  device = "png", dpi = "print", width = 5, height = 5, units = "in"
)
```


## Batch Correction with RUVr

```{r}
library(RUVSeq)
rse <- readRDS(file = "data/rse_reference_no_reps.Rds")

# low_counts <- apply(assay(rse, "counts"), MARGIN = 1, FUN = function(x){all(x < 1000)})
# rse <- rse[-which(low_counts), ]

celltypes <- rse$celltype
set <- newSeqExpressionSet(assay(rse, "counts"),
  phenoData = data.frame(celltypes, row.names = colnames(rse))
)
design <- model.matrix(~celltypes, data = pData(set))
genes <- rownames(rse)
y <- DGEList(counts = counts(set), group = celltypes) %>%
  calcNormFactors(object = ., method = "TMM") %>%
  estimateGLMCommonDisp(y = ., design = design) %>%
  estimateGLMTagwiseDisp(y = ., design = design)

fit <- glmFit(y = y, design = design)
res <- residuals(fit, type = "pearson")

set0 <- RUVr(set, genes, k = 68, res)

ruv <- SummarizedExperiment(
  assays = list(counts = set0@assayData$normalizedCounts),
  rowData = rowData(rse), colData = colData(rse)
)
saveRDS(ruv, file = "data/ruv_reference.Rds")
```

```{r}
library(RUVSeq)
rse <- readRDS(file = "data/rse_reference_noRBC.Rds")

# low_counts <- apply(assay(rse, "counts"), MARGIN = 1, FUN = function(x){all(x < 1000)})
# rse <- rse[-which(low_counts), ]

celltypes <- rse$celltype
celltypes <- droplevels(celltypes)
set <- newSeqExpressionSet(assay(rse, "counts"),
  phenoData = data.frame(celltypes, row.names = colnames(rse))
)
design <- model.matrix(~celltypes, data = pData(set))
genes <- rownames(rse)
y <- DGEList(counts = counts(set), group = celltypes) %>%
  calcNormFactors(object = ., method = "TMM") %>%
  estimateGLMCommonDisp(y = ., design = design) %>%
  estimateGLMTagwiseDisp(y = ., design = design)

fit <- glmFit(y = y, design = design)
res <- residuals(fit, type = "pearson")

set0 <- RUVr(set, genes, k = 67, res)

ruv <- SummarizedExperiment(
  assays = list(counts = set0@assayData$normalizedCounts),
  rowData = rowData(rse), colData = colData(rse)
)
saveRDS(ruv, file = "data/ruv_reference_noRBC.Rds")
```

## CIBERSORT analysis preparation

```{r main cibersort}
library(tidyverse)
library(SummarizedExperiment)
library(DESeq2)

## cibersort takes in a reference expression set to deconvolute a mixture expression set
## read in the reference set
rse <- readRDS(file = "data/ruv_reference.Rds")

rse <- rse[-which(duplicated(rowData(rse)$gene_name)), ]

## read in the mixture set
gtex <- readRDS(file = "data/rse_coronary_recount3_for_cibersort.Rds")

gtex <- gtex[-which(duplicated(rowData(gtex)$gene_name)), ]

## deseq requires the first assay to be counts
## recount3 assigns raw counts to the first slot, so we
## remove raw counts assay from mixture data leaving only the transformed counts

## identify the indices of celltypes of each sample in the reference set
celltypesUnique <- unique(rse$celltype)
celltypes <- rse$celltype

## cibersort requires a phenotype matrix as an input, nCelltypes by nSamples
## the required format uses the value 1 in entry i j to indicate that
## sample j is of celltype i, otherwise the entry takes on the value of 2
## create phenotype identification matrix
classMat <- matrix(
  data = rep.int(x = 2, times = length(celltypesUnique) * ncol(rse)),
  nrow = length(celltypesUnique), ncol = ncol(rse)
)
colnames(classMat) <- colnames(rse)

## mark positions of each celltype in phenotype matrix with 1
for (i in 1:length(celltypesUnique)) {
  tmpInd <- which(celltypes == celltypesUnique[i])
  classMat[i, tmpInd] <- 1
}

## reformat the phenotype matrix according to cibersort specifications
classes <- as.data.frame(classMat) %>%
  mutate(Class = celltypesUnique) %>%
  select(Class, everything())

## normalize the reference set
ref_dds <- DESeqDataSet(rse, design = ~1) %>%
  estimateSizeFactors()

## normalize the mixture set
mix_dds <- DESeqDataSet(gtex, design = ~1) %>%
  estimateSizeFactors()

## transform reference data into the format required by cibersort
## first column gene symbols, remaining columns are normalized count data
reference <- counts(ref_dds, normalized = TRUE) %>%
  round() %>%
  as.data.frame() %>%
  mutate(HGNC = rowData(ref_dds)$gene_name) %>%
  select(HGNC, everything())

## transform mixture data into the format required by cibersort
## first column gene symbols, remaining columns are normalized count data
mixture <- counts(mix_dds, normalized = TRUE) %>%
  round() %>%
  as.data.frame() %>%
  mutate(HGNC = rowData(mix_dds)$gene_name) %>%
  select(HGNC, everything())

## write final tables to input into cibersort
write.table(
  x = classes,
  file = "cibersort/input/ciber_phe_noReps.txt",
  quote = FALSE,
  sep = "\t",
  row.names = FALSE,
  col.names = FALSE
)

write.table(
  x = reference,
  file = "cibersort/input/ciber_ref_noReps.txt",
  quote = FALSE,
  sep = "\t",
  row.names = FALSE,
  col.names = TRUE
)

write.table(
  x = mixture,
  file = "cibersort/input/ciber_mix_noReps.txt",
  quote = FALSE,
  sep = "\t",
  row.names = FALSE,
  col.names = TRUE
)
```

```{r cibersort without rbc}
library(tidyverse)
library(SummarizedExperiment)
library(DESeq2)

## cibersort takes in a reference expression set to deconvolute a mixture expression set
## read in the reference set
rse <- readRDS(file = "data/ruv_reference_noRBC.Rds")

## read in the mixture set
gtex <- readRDS(file = "data/rse_coronary_recount3_for_cibersort.Rds")

## deseq requires the first assay to be counts
## recount3 assigns raw counts to the first slot, so we
## remove raw counts assay from mixture data leaving only the transformed counts

## identify the indices of celltypes of each sample in the reference set
celltypesUnique <- unique(rse$celltype)
celltypes <- rse$celltype

## cibersort requires a phenotype matrix as an input, nCelltypes by nSamples
## the required format uses the value 1 in entry i j to indicate that
## sample j is of celltype i, otherwise the entry takes on the value of 2
## create phenotype identification matrix
classMat <- matrix(
  data = rep.int(x = 2, times = length(celltypes) * ncol(rse)),
  nrow = length(celltypes), ncol = ncol(rse)
)
colnames(classMat) <- colnames(rse)

## mark positions of each celltype in phenotype matrix with 1
for (i in 1:length(celltypesUnique)) {
  tmpInd <- which(celltypes == celltypesUnique[i])
  classMat[i, tmpInd] <- 1
}

## reformat the phenotype matrix according to cibersort specifications
classes <- as.data.frame(classMat) %>%
  mutate(Class = celltypesUnique) %>%
  select(Class, everything())

## normalize the reference set
ref_dds <- DESeqDataSet(rse, design = ~1) %>%
  estimateSizeFactors()

## normalize the mixture set
mix_dds <- DESeqDataSet(gtex, design = ~1) %>%
  estimateSizeFactors()

## transform reference data into the format required by cibersort
## first column gene symbols, remaining columns are normalized count data
reference <- counts(ref_dds, normalized = TRUE) %>%
  round() %>%
  as.data.frame() %>%
  mutate(HGNC = rowData(ref_dds)$gene_name) %>%
  select(HGNC, everything())

## transform mixture data into the format required by cibersort
## first column gene symbols, remaining columns are normalized count data
mixture <- counts(mix_dds, normalized = TRUE) %>%
  round() %>%
  as.data.frame() %>%
  mutate(HGNC = rowData(mix_dds)$gene_name) %>%
  select(HGNC, everything())

## write final tables to input into cibersort
write.table(
  x = classes,
  file = "cibersort/input/ciber_phe_recount3_noBlood.txt",
  quote = FALSE,
  sep = "\t",
  row.names = FALSE,
  col.names = FALSE
)

write.table(
  x = reference,
  file = "cibersort/input/ciber_ref_recount3_noBlood.txt",
  quote = FALSE,
  sep = "\t",
  row.names = FALSE,
  col.names = TRUE
)
```

## CIBERSORT review

```{r}
gtex_n235 <- readRDS(file = "data/gtex_n235_exp_scores.Rds")
ciberOut <- read.table(file = "cibersort/output/final.txt", header = TRUE)

ciberOut <- data.frame(ciberOut[, 1:8])

ciberOut$Mixture <- factor(gtex_n235$histologyID)
colnames(ciberOut) <- c("Sample", "CMC", "END", "LYM", "SMC", "MAC", "RBC", "ADI")

ciberOut$Sample <- fct_reorder2(ciberOut$Sample, ciberOut$ADI, ciberOut$SMC)
ciberDf <- ciberOut
ciberDf <- pivot_longer(ciberDf, cols = 2:8, names_to = "Cell type", values_to = "Proportion")

ciberDf$`Cell type` <- factor(ciberDf$`Cell type`,
  levels = c(
    "ADI",
    "CMC",
    "MAC",
    "LYM",
    "END",
    "SMC",
    "RBC"
  )
)

gg_ciber <- ggplot(
  ciberDf,
  aes(x = Sample, y = Proportion, fill = `Cell type`, group = `Cell type`)
) +
  geom_bar(stat = "identity", width = 1) +
  scale_fill_mnmlab(palette = "martink8") +
  labs( # title = "Reference-Based Deconvolution of Plasma miRNA with CIBERSORT",
    x = "Sample",
    y = "Estimated % of Tissue",
    fill = "Cell type"
  ) +
  theme(
    panel.background = element_rect(fill = "white"),
    axis.text.x = element_blank(),
    axis.ticks.x = element_blank(),
    panel.grid = element_blank(),
    legend.title = element_text(size = 11),
    legend.text = element_text(size = 11),
    legend.key.size = unit(2, "line")
  )
ggsave(
  filename = "figures/gg_ciber.png", plot = gg_ciber,
  device = "png", dpi = "print", width = 5, height = 5, units = "in"
)

gtex_n235$ADI <- ciberOut$ADI
gtex_n235$CMC <- ciberOut$CMC
gtex_n235$SMC <- ciberOut$SMC
gtex_n235$END <- ciberOut$END
gtex_n235$RBC <- ciberOut$RBC
gtex_n235$LYM <- ciberOut$LYM
gtex_n235$MAC <- ciberOut$MAC

gtex_n235$ADIz <- (gtex_n235$ADI - mean(gtex_n235$ADI)) / sd(gtex_n235$ADI)
gtex_n235$CMCz <- (gtex_n235$CMC - mean(gtex_n235$CMC)) / sd(gtex_n235$CMC)
gtex_n235$SMCz <- (gtex_n235$SMC - mean(gtex_n235$SMC)) / sd(gtex_n235$SMC)
gtex_n235$ENDz <- (gtex_n235$END - mean(gtex_n235$END)) / sd(gtex_n235$END)
gtex_n235$RBCz <- (gtex_n235$RBC - mean(gtex_n235$RBC)) / sd(gtex_n235$RBC)
gtex_n235$LYMz <- (gtex_n235$LYM - mean(gtex_n235$LYM)) / sd(gtex_n235$LYM)
gtex_n235$MACz <- (gtex_n235$MAC - mean(gtex_n235$MAC)) / sd(gtex_n235$MAC)

gtex_n138 <- gtex_n235[, c(subsetInd01, subsetInd45)]

saveRDS(gtex_n138, file = "data/gtex_n138_for_deseq.Rds")

plotDf <- data.frame(
  Celltype = rep(c("SMC", "END", "MAC", "LYM", "ADI", "CMC"), each = 138),
  CIB = c(
    gtex_n138$SMC,
    gtex_n138$END,
    gtex_n138$MAC,
    gtex_n138$LYM,
    gtex_n138$ADI,
    gtex_n138$CMC
  ),
  CibScore = c(
    gtex_n138$SMCz,
    gtex_n138$ENDz,
    gtex_n138$MACz,
    gtex_n138$LYMz,
    gtex_n138$ADIz,
    gtex_n138$CMCz
  ),
  ExpScore = c(
    gtex_n138$SmcScore,
    gtex_n138$EndScore,
    gtex_n138$MacScore,
    gtex_n138$LymScore,
    gtex_n138$AdiScore,
    gtex_n138$CmcScore
  ),
  Plaque = rep(gtex_n138$plaque, times = 6),
  Sample = rep(gtex_n138$histologyID, times = 6)
)

plotDf$Plaque <- factor(plotDf$Plaque, levels = c(0, 1), labels = c("No", "Yes"))

gg_cib <- ggplot(plotDf, aes(x = Sample, y = CIB, fill = Celltype, group = Celltype)) +
  geom_bar(stat = "identity")

gg_cib_exp_comparison <- ggplot(plotDf, aes(x = CibScore, y = ExpScore, color = Plaque)) +
  geom_point(alpha = 0.5) +
  facet_wrap(~Celltype, scales = "free") +
  labs( # title = "CIBERSORT estimates against gene expression per celltype",
    x = "CIBERSORT (centered and scaled proportion estimates)",
    y = "Expression (centered and scaled VST counts)"
  ) +
  scale_color_manual(values = c("No" = "#004488", "Yes" = "#BB5566"))

ggsave(
  filename = "figures/cib_exp_comparison.png",
  plot = gg_cib_exp_comparison,
  device = "png",
  dpi = "print",
  width = 5,
  height = 5,
  units = "in"
)
```

## Power Calculations

```{r}
library(PROPER)

simOpts1 <- RNAseq.SimOptions.2grp(
  ngenes = 12642, p.DE = 0.01, lOD = "cheung",
  lBaselineExpr = "cheung"
)
simOpts5 <- RNAseq.SimOptions.2grp(
  ngenes = 12642, p.DE = 0.05, lOD = "cheung",
  lBaselineExpr = "cheung"
)
simOpts10 <- RNAseq.SimOptions.2grp(
  ngenes = 12642, p.DE = 0.1, lOD = "cheung",
  lBaselineExpr = "cheung"
)

simOptsList <- list(simOpts1, simOpts5, simOpts10)

simCheung <- function(opts) {
  runSims(
    Nreps = 53,
    Nreps2 = 85,
    sim.opts = opts,
    DEmethod = "DESeq2",
    nsims = 20
  )
}

simResCheung <- lapply(simOptsList, simCheung)

powers_lfc585 <- lapply(simResCheung,
  comparePower,
  alpha.type = "fdr",
  alpha.nominal = 0.1,
  stratify.by = "dispersion",
  target.by = "lfc",
  delta = 0.585
)

powers_lfc1 <- lapply(simResCheung,
  comparePower,
  alpha.type = "fdr",
  alpha.nominal = 0.1,
  stratify.by = "dispersion",
  target.by = "lfc",
  delta = 1
)

powers_eff585 <- lapply(simResCheung,
  comparePower,
  alpha.type = "fdr",
  alpha.nominal = 0.1,
  stratify.by = "dispersion",
  target.by = "effectsize",
  delta = 0.585
)

powers_eff1 <- lapply(simResCheung,
  comparePower,
  alpha.type = "fdr",
  alpha.nominal = 0.1,
  stratify.by = "dispersion",
  target.by = "effectsize",
  delta = 1
)

powersList <- list(
  summaryPower(powers_lfc585[[1]]),
  summaryPower(powers_lfc585[[2]]),
  summaryPower(powers_lfc585[[3]]),
  summaryPower(powers_lfc1[[1]]),
  summaryPower(powers_lfc1[[2]]),
  summaryPower(powers_lfc1[[3]]),
  summaryPower(powers_eff585[[1]]),
  summaryPower(powers_eff585[[2]]),
  summaryPower(powers_eff585[[3]]),
  summaryPower(powers_eff1[[1]]),
  summaryPower(powers_eff1[[2]]),
  summaryPower(powers_eff1[[3]])
)

pDE <- rep(c(1, 5, 10), 4)

targetDelta <- c(rep("lfc = 0.585", 3), rep("lfc = 1", 3), rep("eff = 0.585", 3), rep("eff = 1", 3))

powerDf <- do.call(rbind, powersList) %>%
  data.frame() %>%
  mutate("Percent DE" = pDE, "Target" = targetDelta) %>%
  select("Target", "Percent DE", everything()) %>%
  select(-"FDC")

colnames(powerDf) <- c(
  "Target",
  "PercentDE",
  "SS1",
  "SS2",
  "Nominal FDR",
  "Actual FDR",
  "Marginal Power",
  "Avg TD",
  "Avg FD"
)
```

## DESeq2 analysis

```{r}
deseq_noComp <- DESeqDataSet(gtex_n138, design = ~ plaque + sex + age + year) %>% DESeq()

deseq_expScores <- DESeqDataSet(gtex_n138, design = ~ plaque + sex + age + year + SmcScore + CmcScore + RbcScore) %>% DESeq()

deseq_cibScores <- DESeqDataSet(gtex_n138, design = ~ plaque + sex + age + year + SMC + CMC + RbcScore) %>% DESeq()

res_noComp <- reduceResults(deseq_noComp[-which(rowData(gtex_n138)$gene_name %in% markerGenesVec), ], contrast = c("plaque", 1, 0))

res_noComp_genes <- res_noComp$HGNC[order(abs(res_noComp$log2FoldChange), decreasing = TRUE)][1:200]

res_expScores <- reduceResults(deseq_expScores[-which(rowData(gtex_n138)$gene_name %in% markerGenesVec), ], contrast = c("plaque", 1, 0))

res_exp_genes <- res_expScores$HGNC[order(abs(res_expScores$log2FoldChange), decreasing = TRUE)][1:400]

res_cibScores <- reduceResults(deseq_cibScores[-which(rowData(gtex_n138)$gene_name %in% markerGenesVec), ], contrast = c("plaque", 1, 0))

res_cib_genes <- res_cibScores$HGNC[order(abs(res_cibScores$log2FoldChange), decreasing = TRUE)][1:400]

full_res_noComp <- data.frame(results(deseq_noComp[-which(rowData(gtex_n138)$gene_name %in% markerGenesVec), ], contrast = c("plaque", 1, 0))) %>% mutate(HGNC = rowData(gtex_n138)$gene_name[-which(rowData(gtex_n138)$gene_name %in% markerGenesVec)])

full_res_expScores <- data.frame(results(deseq_expScores[-which(rowData(gtex_n138)$gene_name %in% markerGenesVec), ], contrast = c("plaque", 1, 0))) %>% mutate(HGNC = rowData(gtex_n138)$gene_name[-which(rowData(gtex_n138)$gene_name %in% markerGenesVec)])

full_res_cibScores <- data.frame(results(deseq_cibScores[-which(rowData(gtex_n138)$gene_name %in% markerGenesVec), ], contrast = c("plaque", 1, 0))) %>% mutate(HGNC = rowData(gtex_n138)$gene_name[-which(rowData(gtex_n138)$gene_name %in% markerGenesVec)])

sig_genes <- union(res_cib_genes, res_exp_genes) %>% union(., res_noComp_genes)

sig_genes <- sig_genes[-which(sig_genes %in% igGenes)]

set.seed(02241950)

sig_genes <- sample(sig_genes, 100)

pDF <- data.frame(
  Gene = common_genes,
  p1 = vector(length = length(common_genes)),
  p2 = vector(length = length(common_genes)),
  eff1 = vector(length = length(common_genes)),
  eff2 = vector(length = length(common_genes))
)

for (i in 1:length(common_genes)) {
  ind1 <- which(full_res_cibScores$HGNC == common_genes[i])
  ind2 <- which(full_res_expScores$HGNC == common_genes[i])

  pDF$pCIB[i] <- full_res_cibScores$padj[ind1]
  pDF$effCIB[i] <- full_res_cibScores$log2FoldChange[ind1]
  pDF$pEXP[i] <- full_res_expScores$padj[ind2]
  pDF$effEXP[i] <- full_res_expScores$log2FoldChange[ind2]
}

gg_p <- ggplot(pDF, aes(x = effCIB - effEXP, y = log10(pEXP) - log10(pCIB))) +
  geom_point(alpha = 0.2) +
  labs( # title = "Comparison of DESeq Results depending on composition estimate method",
    x = TeX("$l2fc_{cib} - l2fc_{exp}$"),
    y = TeX("$\\log_{10}padj_{exp} - \\log_{10}padj_{cib}$")
  )
ggsave(
  filename = "figures/gg_p_v_e.png", plot = gg_p,
  device = "png", dpi = "print", width = 5, height = 5, units = "in"
)

indNonU <- which(sig_genes %in% res_noComp$HGNC[res_noComp$log2FoldChange >= 0])
indNonD <- which(sig_genes %in% res_noComp$HGNC[res_noComp$log2FoldChange < 0])
sigNon <- rep(0, length = length(sig_genes))
sigNon[indNonD] <- 1
sigNon[indNonU] <- 2
effNon <- rep(0, length = length(sig_genes))
for (i in 1:length(sig_genes)) {
  tmpInd <- which(full_res_noComp$HGNC == sig_genes[i])
  effNon[i] <- full_res_noComp$log2FoldChange[tmpInd] %>% round(digits = 3)
}

indcibU <- which(sig_genes %in% res_cibScores$HGNC[res_cibScores$log2FoldChange >= 0])
indcibD <- which(sig_genes %in% res_cibScores$HGNC[res_cibScores$log2FoldChange < 0])
sigcib <- rep(0, length = length(sig_genes))
sigcib[indcibD] <- 1
sigcib[indcibU] <- 2
effcib <- rep(0, length = length(sig_genes))
for (i in 1:length(sig_genes)) {
  tmpInd <- which(full_res_cibScores$HGNC == sig_genes[i])
  effcib[i] <- full_res_cibScores$log2FoldChange[tmpInd] %>% round(digits = 3)
}

indexpU <- which(sig_genes %in% res_expScores$HGNC[res_expScores$log2FoldChange >= 0])
indexpD <- which(sig_genes %in% res_expScores$HGNC[res_expScores$log2FoldChange < 0])
sigexp <- rep(0, length = length(sig_genes))
sigexp[indexpD] <- 1
sigexp[indexpU] <- 2
effexp <- rep(0, length = length(sig_genes))
for (i in 1:length(sig_genes)) {
  tmpInd <- which(full_res_expScores$HGNC == sig_genes[i])
  effexp[i] <- full_res_expScores$log2FoldChange[tmpInd] %>% round(digits = 3)
}

df <- data.frame(
  Gene = factor(sig_genes),
  sigNon = sigNon, effNon = effNon,
  sigCib = sigcib, effCib = effcib,
  sigExp = sigexp, effExp = effexp
)

df$Gene <- fct_reorder(df$Gene, df$effNon)

dfLong <- pivot_longer(df, cols = 2:7, names_to = c(".value", "Model"), names_pattern = "(...)(...)")

dfLong$Model <- factor(dfLong$Model,
  labels = c("None", "CIB", "EXP"),
  levels = c("Non", "Cib", "Exp")
)

dfLong$sig[dfLong$sig == 2] <- 1

dfLong$sig <- factor(dfLong$sig, levels = 0:1, labels = c("No", "Yes"))

ggplot(dfLong, aes(x = Gene, y = eff)) +
  geom_quasirandom(aes(color = Model, shape = factor(sig))) +
  geom_boxplot(width = .15, fill = NA, outlier.shape = NA) +
  scale_color_mnmlab(palette = "highContrast") +
  theme_bw() +
  theme(axis.text.x = element_text(angle = 45, hjust = .9, size = 4, face = "italic")) +
  labs(#title = "Distribution of L2FC across Models", 
       y = "L2FC", 
       shape = "Sig.",
       color = "Comp.")

ggsave(
  filename = "figures/gg_lfc.png",
  device = "png", dpi = "print", width = 9, height = 6, units = "in"
)
```
````
