## Supplementary figures and images for "Considerations for Deconvolution: A Case Study with GTEx Coronary Artery Tissues"

### cib_exp_comparison.png

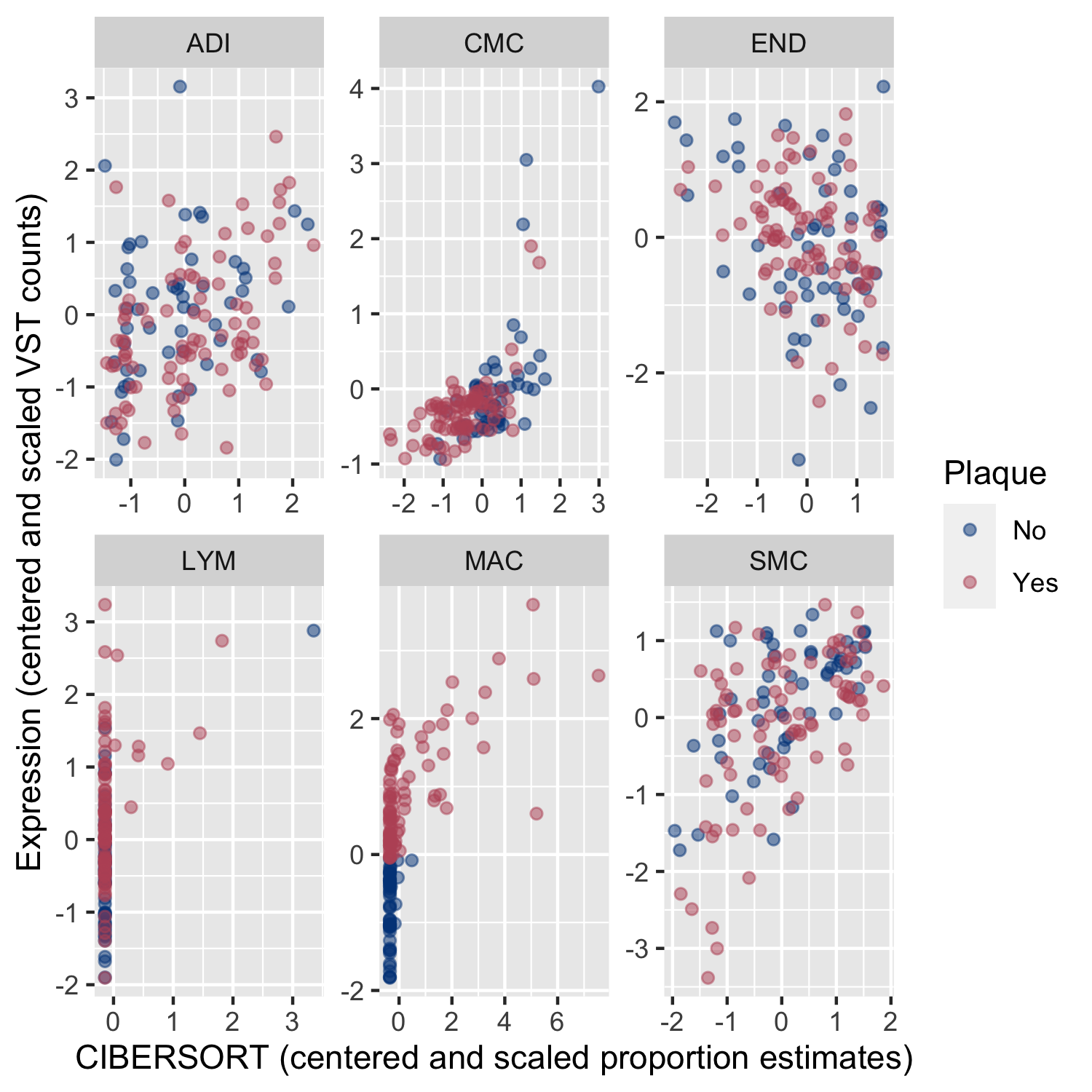

### cibersort_corr.png

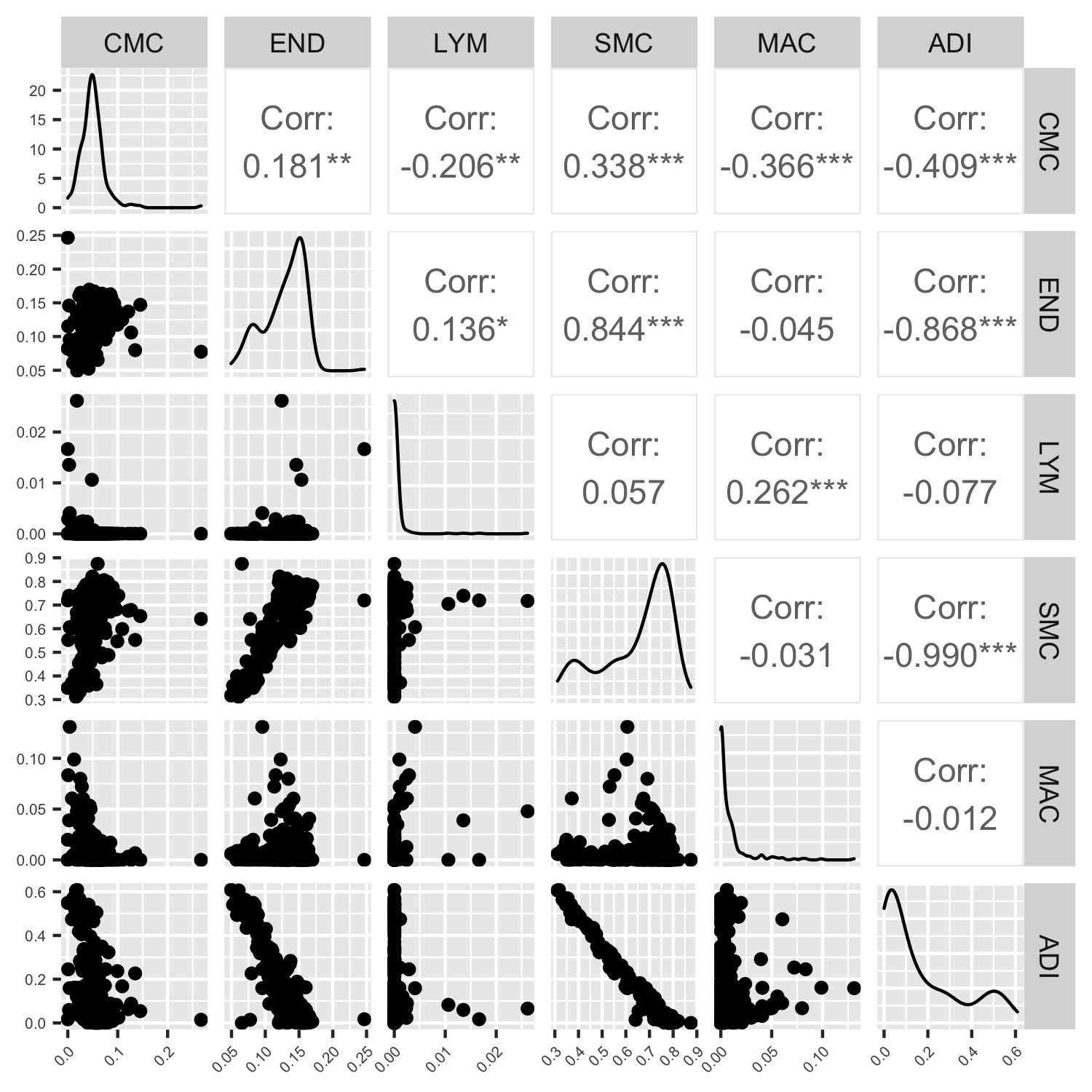

### cibersort_results.png

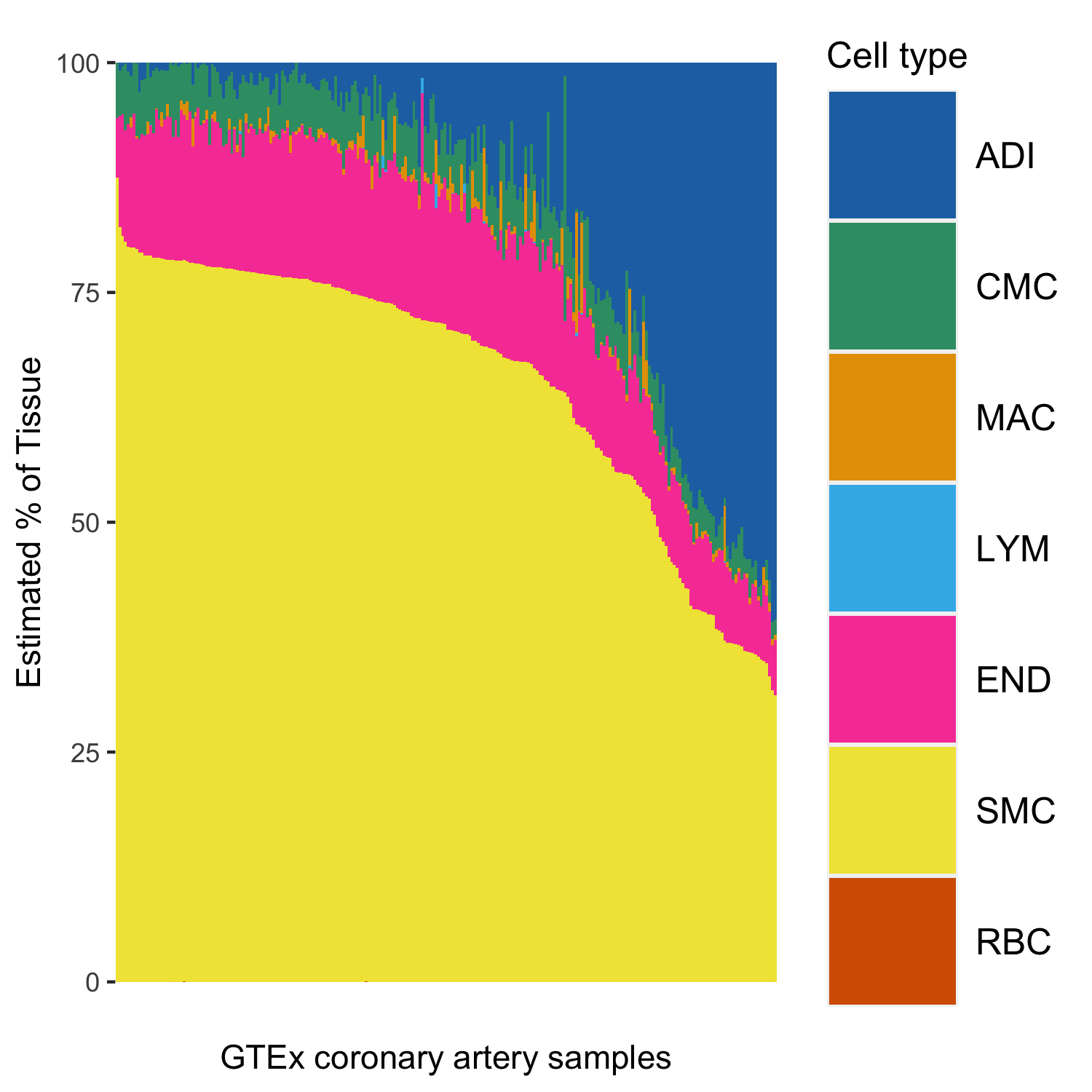

### Fat_vein_tSNE.png

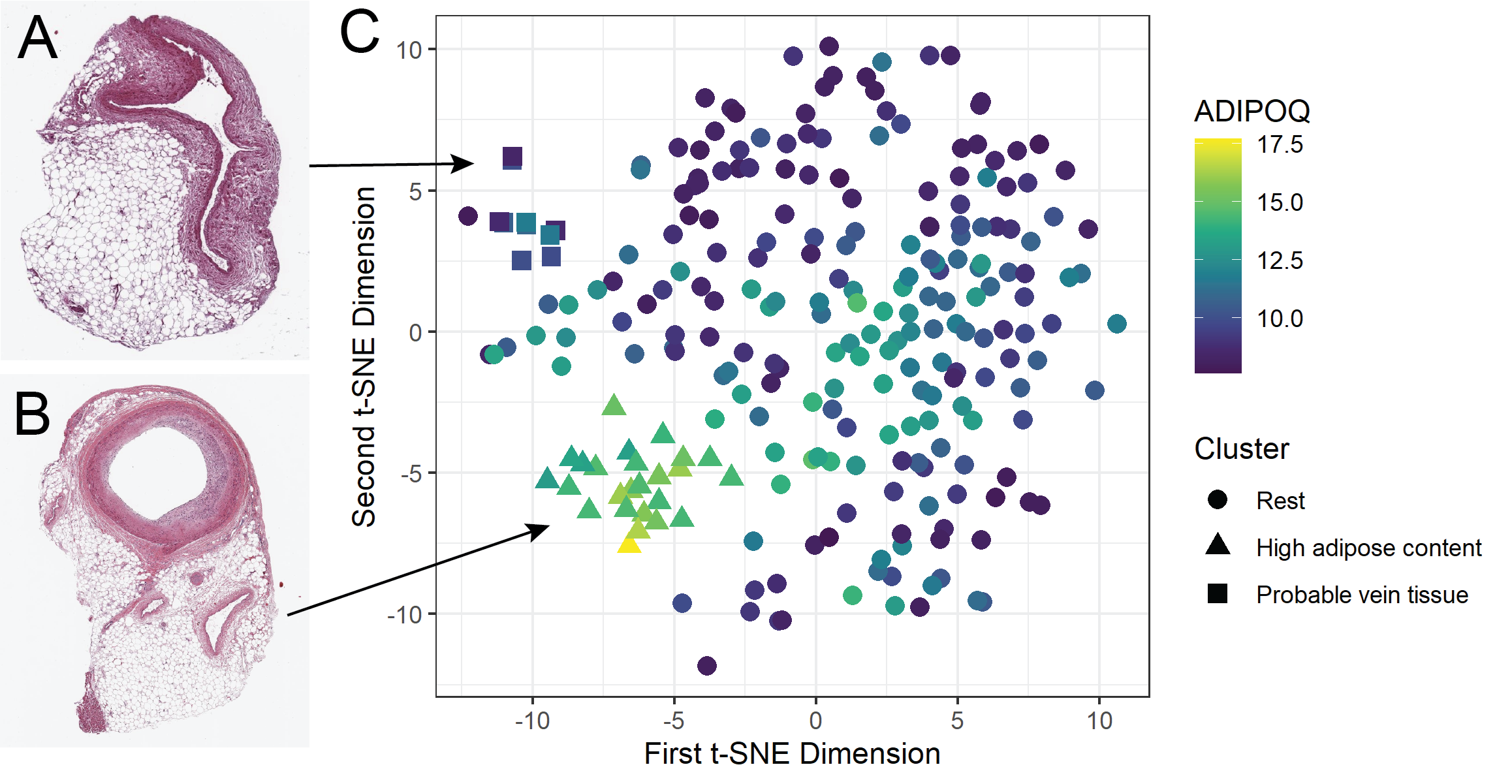

### gg_ciber.png

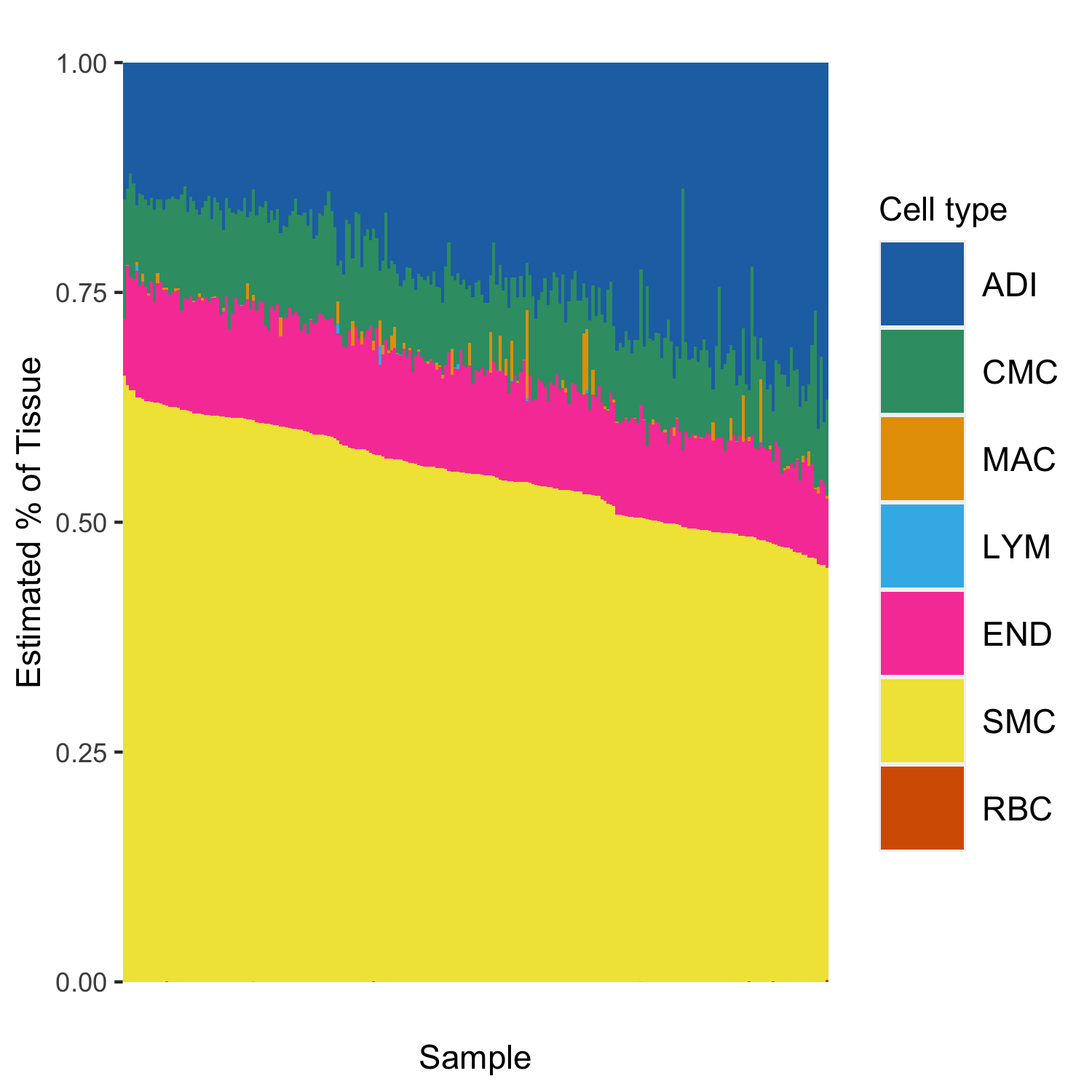

### gg_cmd.png

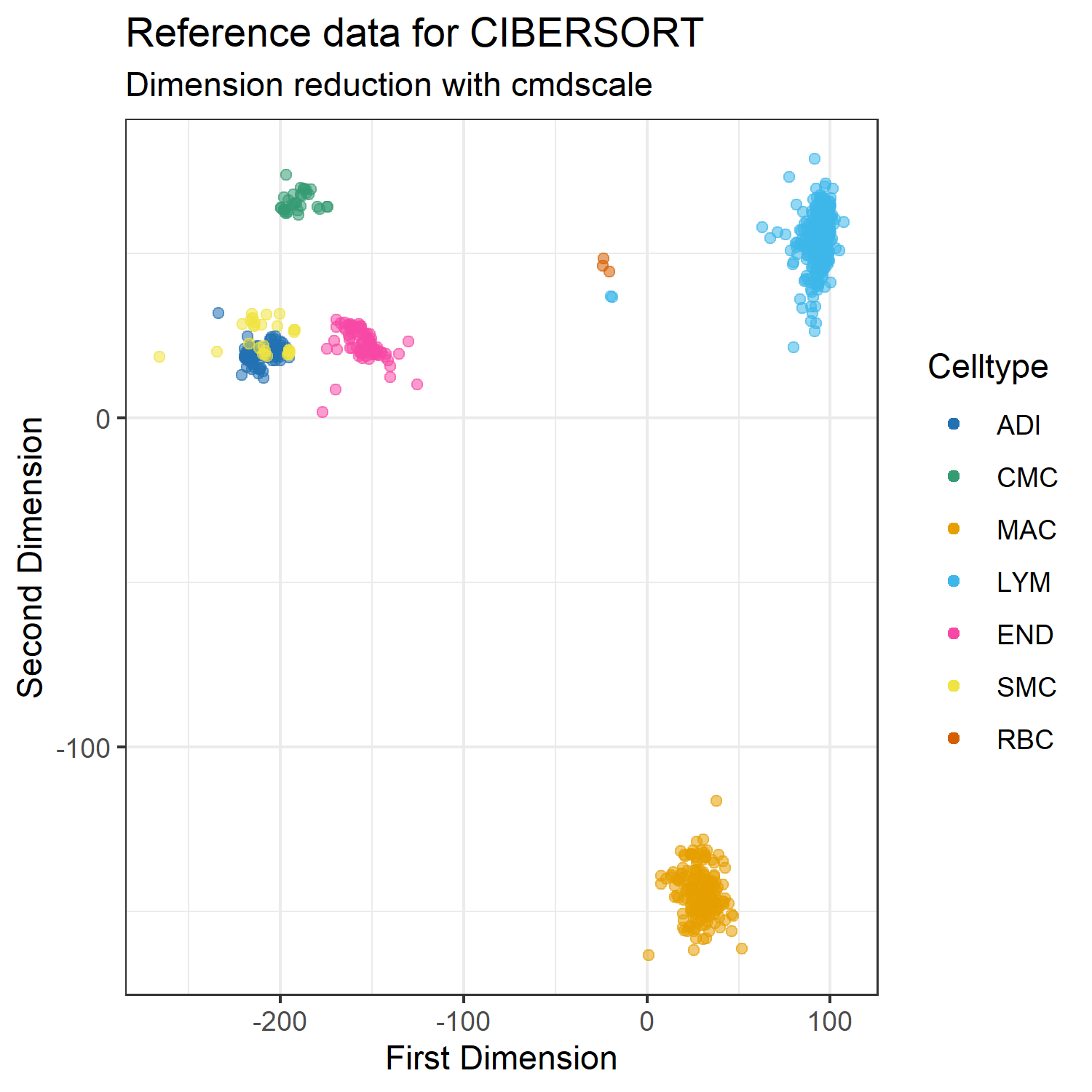

### gg_lfc.png

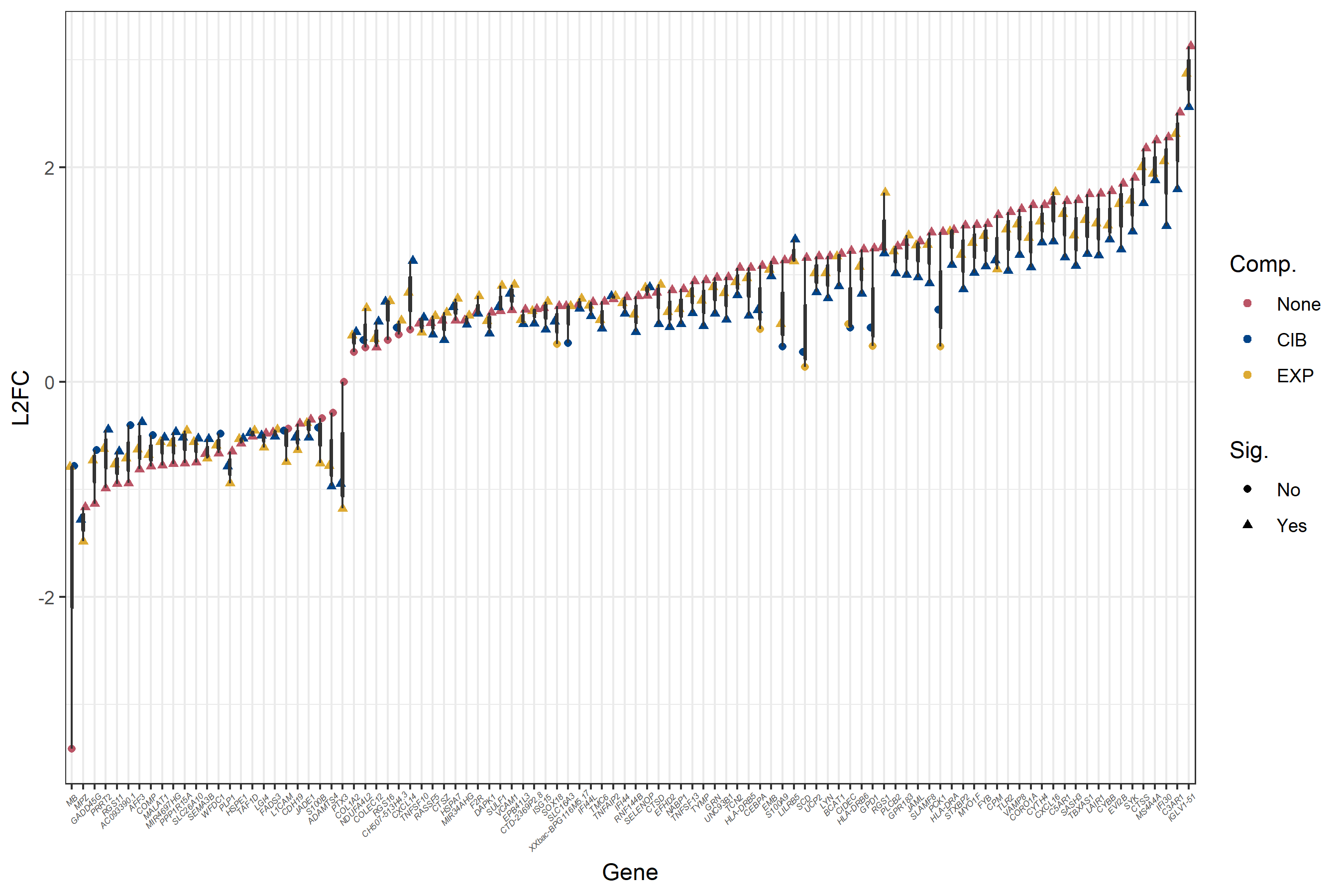

### gg_pca_gtex.png

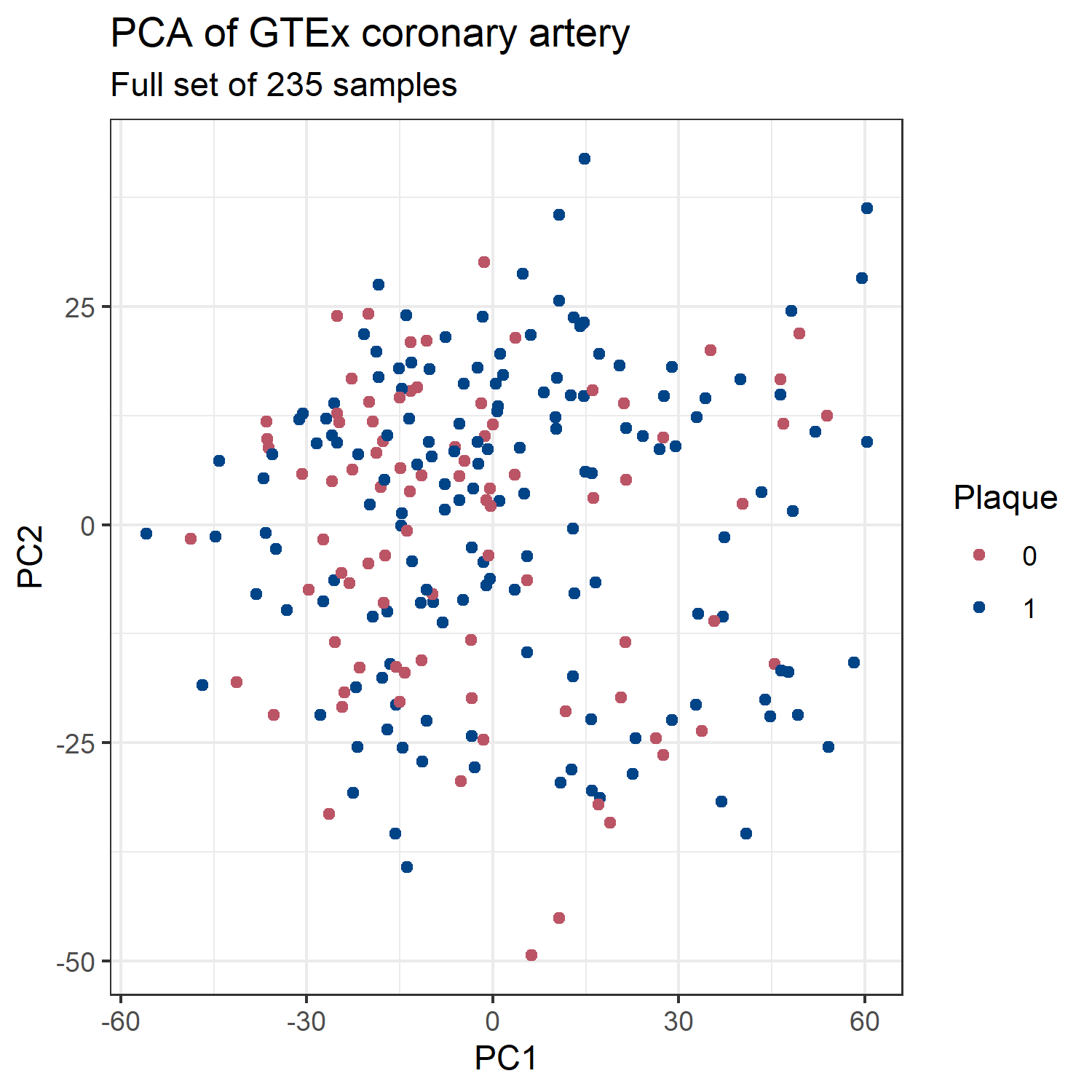

### gg_pca_gtex_top100.png

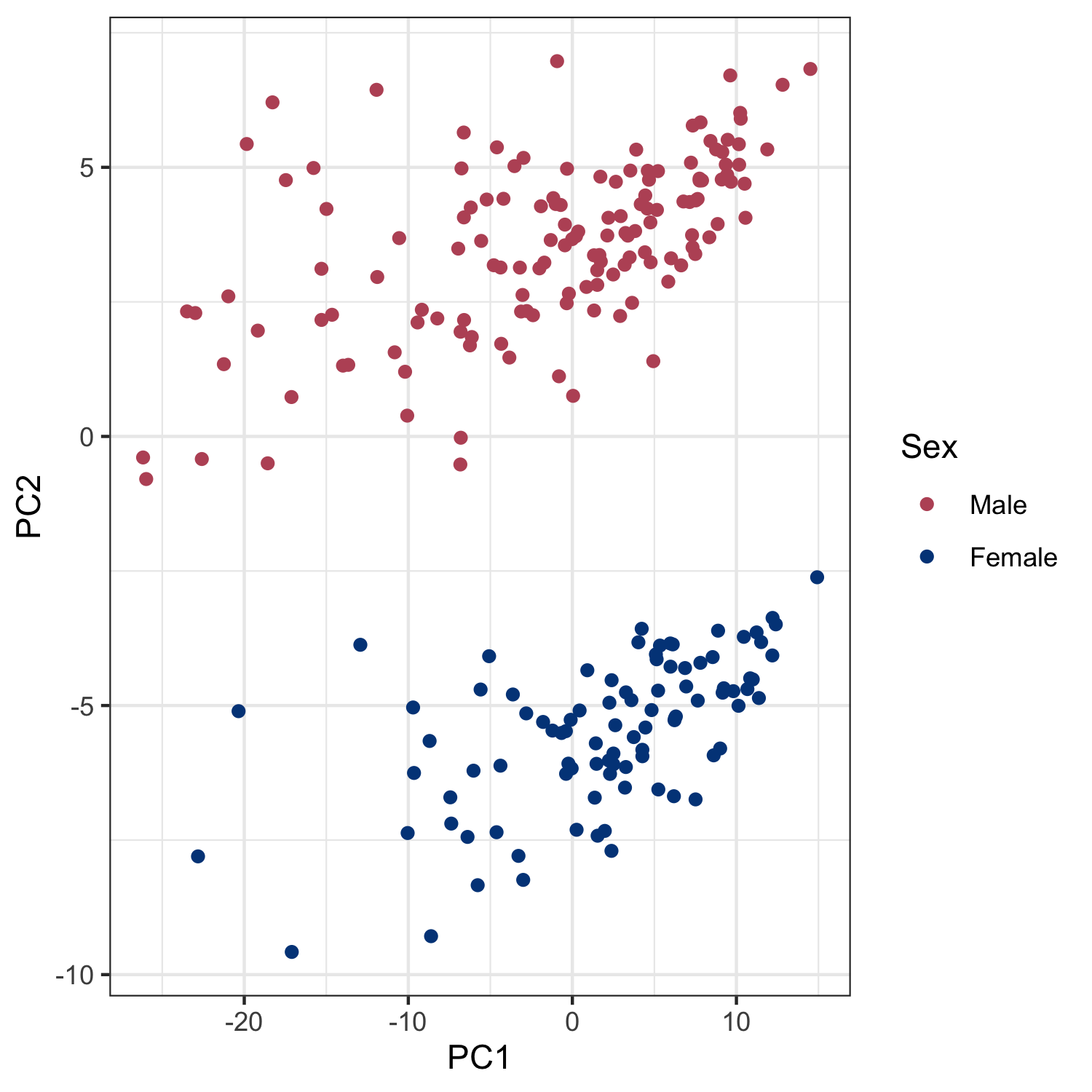

### gg_pca_gtexSub.png

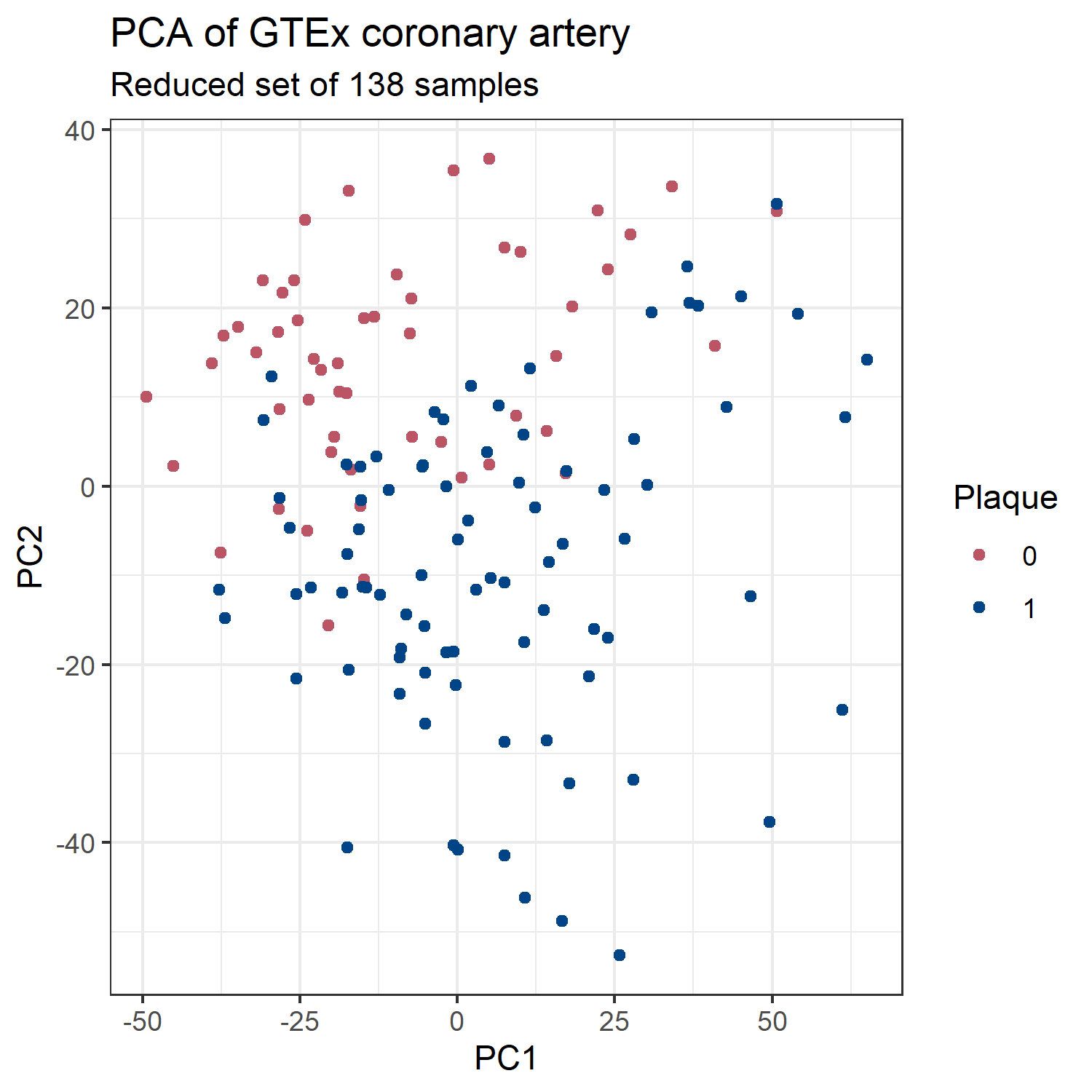

### gg_tsne.png

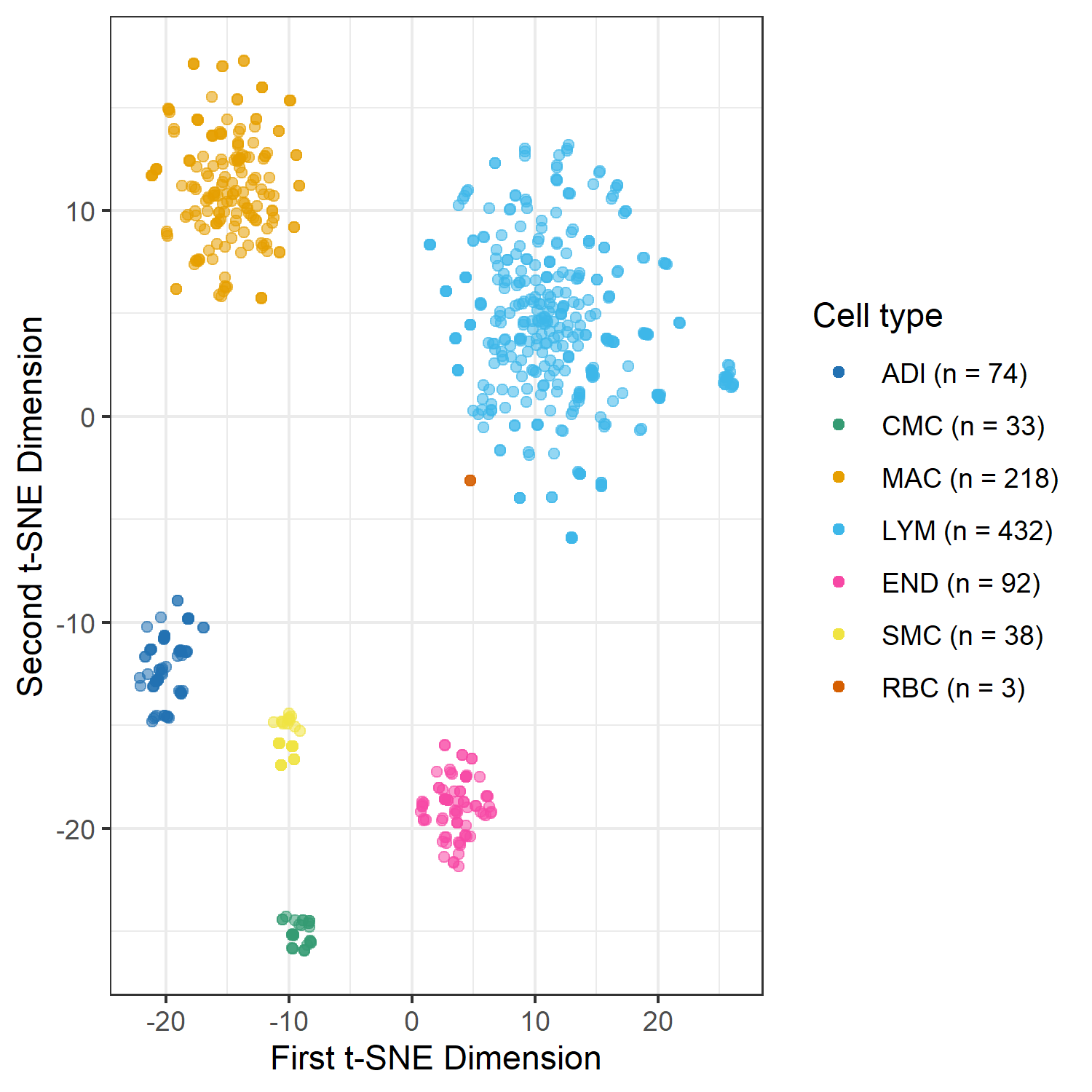

### gg_umap.png

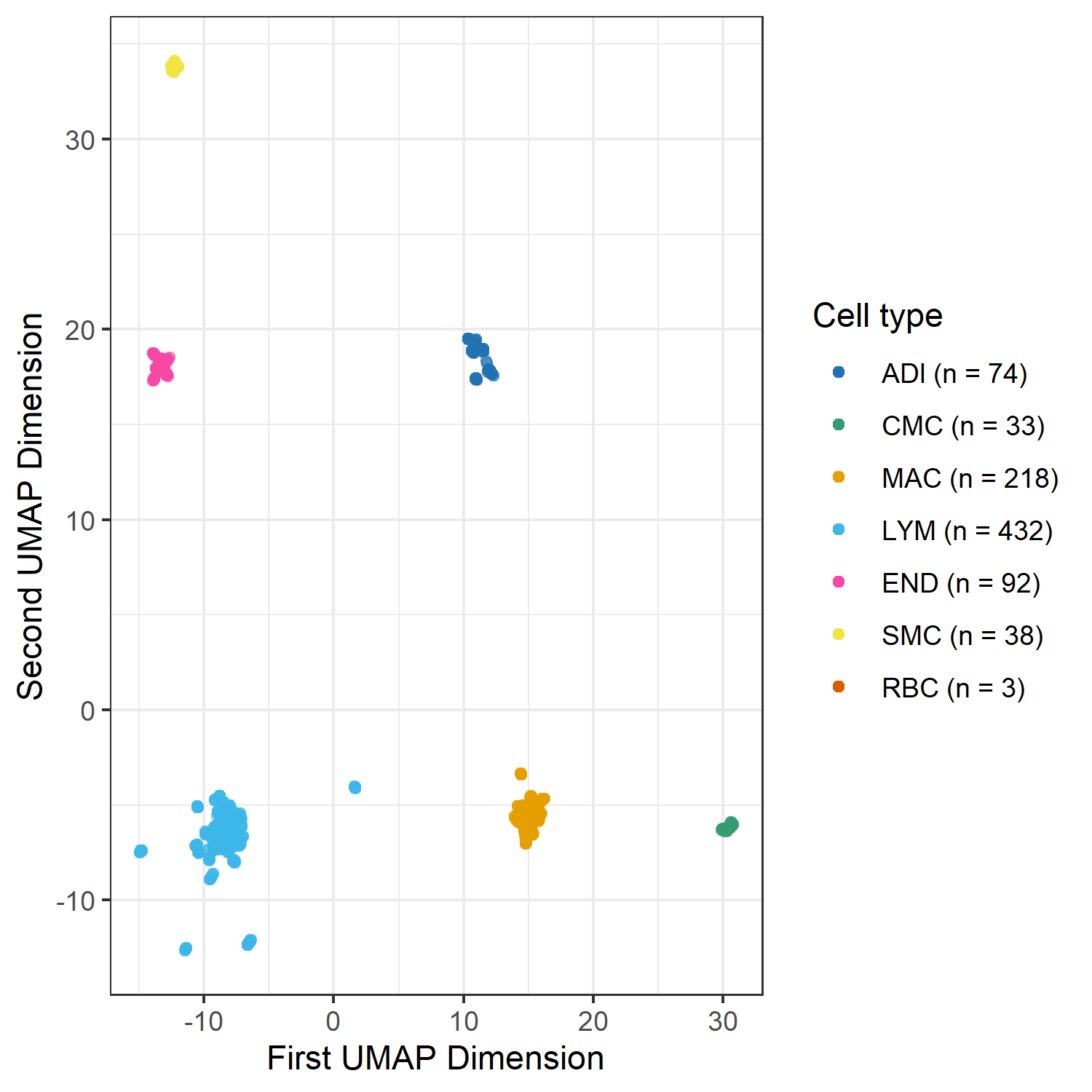

### noBloodComparison.png

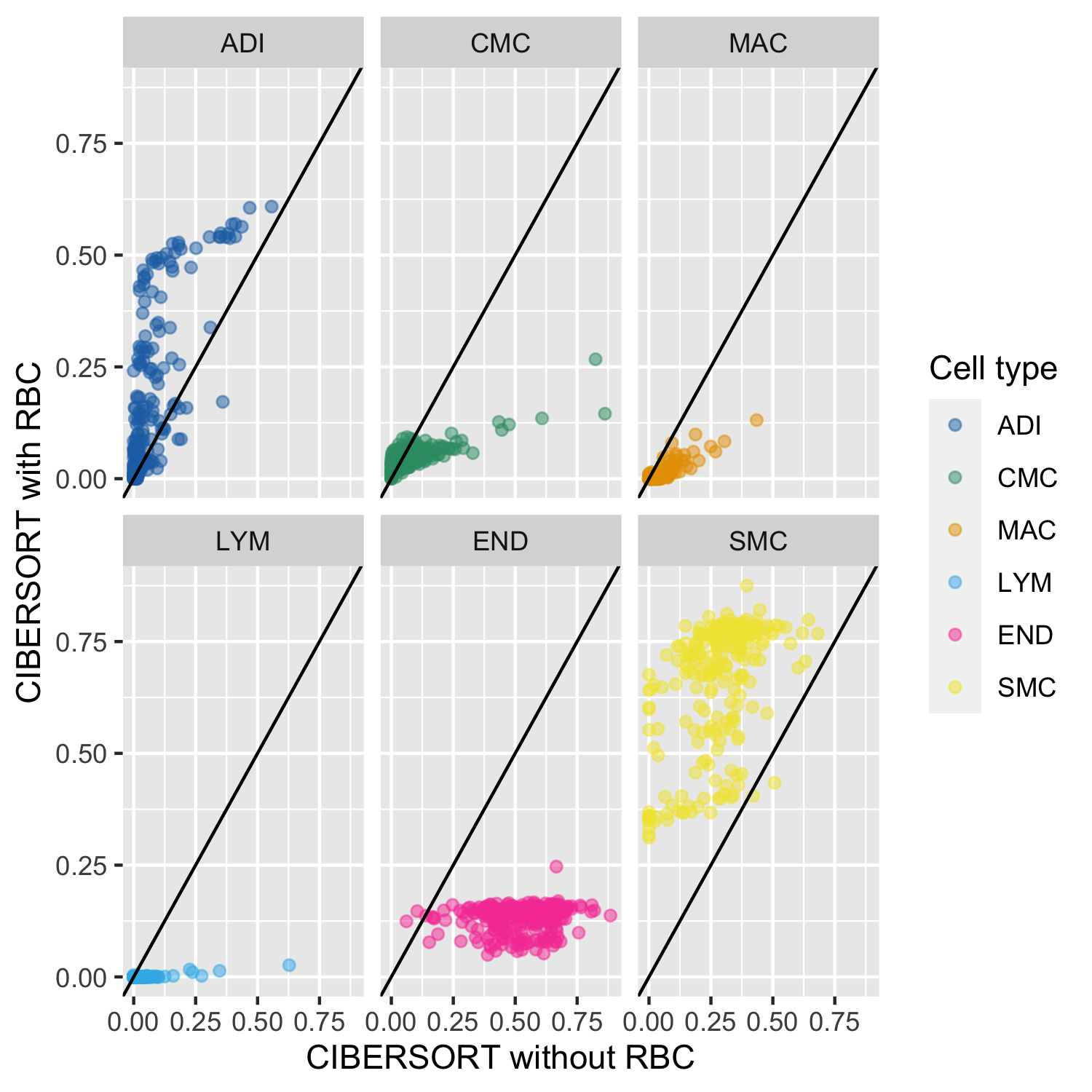

### tsne_vein_clusters.png

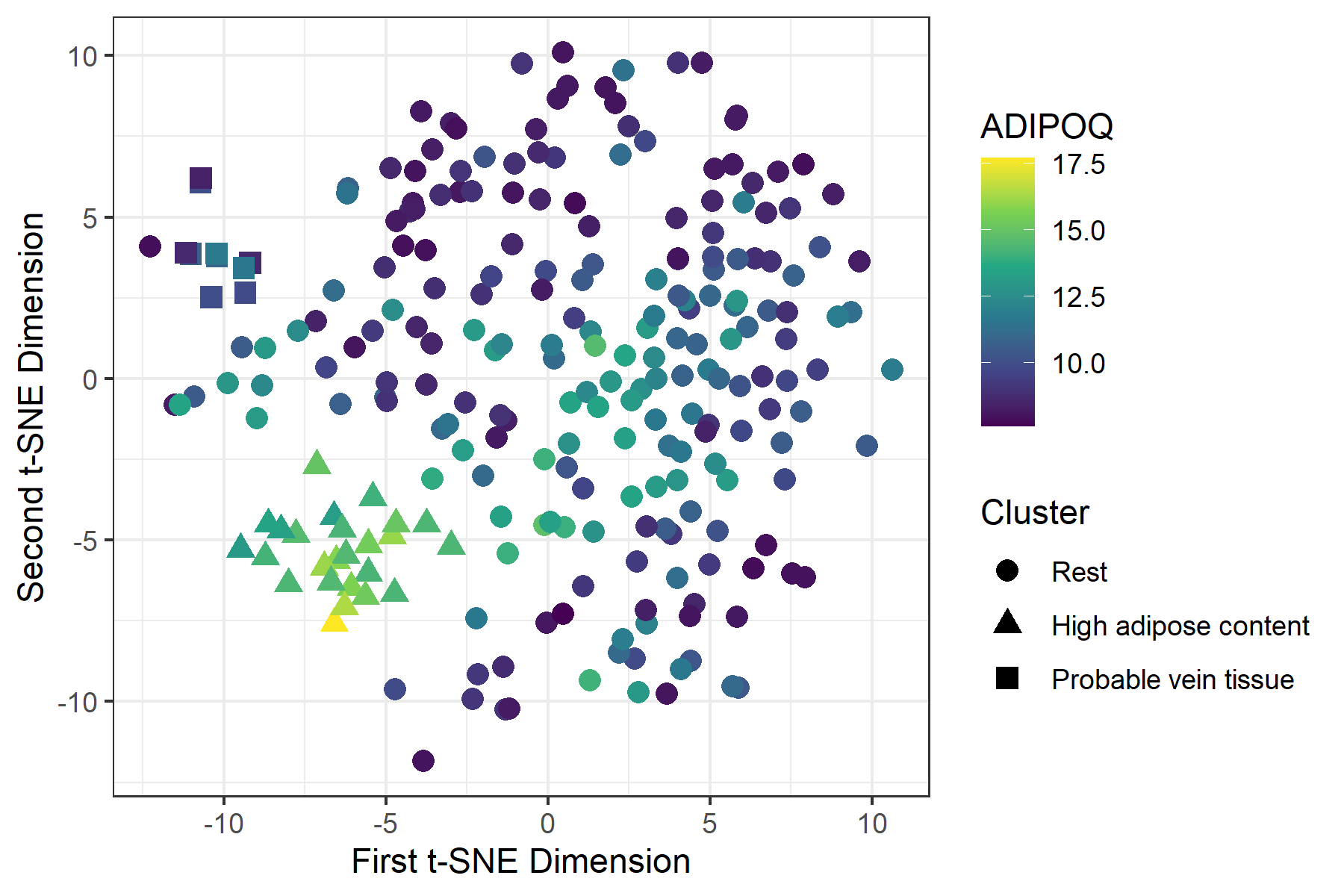
